# Multiplex, single-cell CRISPRa screening for cell type specific regulatory elements

**DOI:** 10.1101/2023.03.28.534017

**Authors:** Florence M. Chardon, Troy A. McDiarmid, Nicholas F. Page, Riza M. Daza, Beth Martin, Silvia Domcke, Samuel G. Regalado, Jean-Benoît Lalanne, Diego Calderon, Xiaoyi Li, Lea M. Starita, Stephan J. Sanders, Nadav Ahituv, Jay Shendure

## Abstract

CRISPR-based gene activation (CRISPRa) is a promising therapeutic approach for gene therapy, upregulating gene expression by targeting promoters or enhancers in a tissue/cell-type specific manner. Here, we describe an experimental framework that combines highly multiplexed perturbations with single-cell RNA sequencing (sc-RNA-seq) to identify cell-type-specific, CRISPRa-responsive *cis-*regulatory elements and the gene(s) they regulate. Random combinations of many gRNAs are introduced to each of many cells, which are then profiled and partitioned into test and control groups to test for effect(s) of CRISPRa perturbations of both enhancers and promoters on the expression of neighboring genes. Applying this method to a library of 493 gRNAs targeting candidate *cis-*regulatory elements in both K562 cells and iPSC-derived excitatory neurons, we identify gRNAs capable of specifically upregulating intended target genes and no other neighboring genes within 1 Mb, including gRNAs yielding upregulation of six autism spectrum disorder (ASD) and neurodevelopmental disorder (NDD) risk genes in neurons. A consistent pattern is that the responsiveness of individual enhancers to CRISPRa is restricted by cell type, implying a dependency on either chromatin landscape and/or additional *trans-*acting factors for successful gene activation. The approach outlined here may facilitate large-scale screens for gRNAs that activate therapeutically relevant genes in a cell type-specific manner.

## Introduction

There are millions of candidate *cis-*regulatory elements (cCREs) in the human genome, yet only a handful have been functionally validated and confidently linked to their target gene(s)^1^. Recently, we and others have combined CRISPR-interference (CRISPRi) and sc-RNA-seq to scalably validate distal cCREs, while also linking them to the gene(s) that they regulate^1–4^. However, to date, the vast majority of work in the field has focused on screening candidate regulatory elements for *necessity*, with only a few studies screening for *sufficiency* in the endogenous context.

CRISPR-activation (CRISPRa) is a versatile approach that allows one to test for the sufficiency of cCRE activity^5–8^. CRISPRa screens of noncoding regulatory elements have at least four potential advantages over CRISPRi screens. First, as noted above, CRISPRa can identify cCREs that are sufficient even if not singularly necessary to drive target gene expression. Second, CRISPRa can identify elements that, when targeted, may upregulate already active genes above their baseline levels. Third, CRISPRa has the potential to discover inactive regions that, when transcriptional activation machinery is recruited to them, can act as active enhancers and increase expression of nearby genes^9^. Finally, CRISPRa has the potential to identify cCRE-targeting gRNAs whose activity is cell type-specific, opening the door to “*cis* regulatory therapy” (CRT) for haploinsufficient and other low-dosage associated disorders, as recently demonstrated for monogenic forms of obesity and autism spectrum disorder^10,11^. However, despite these potential advantages, CRISPRa targeting of noncoding regulatory elements has mostly been deployed in an *ad hoc* manner^9,12–14^, meaning typically focusing on a single target gene, and furthermore in workhorse cancer cell lines rather than more therapeutically relevant *in vitro* models.

Here, we present a scalable framework in which we introduce multiple, random combinations of CRISPRa perturbations to each of many cells followed by sc-RNA-seq (**Fig. 1**), analogous to an approach that we previously developed for CRISPRi screening^2,4^. Computational partitioning of cells into test and control groups based on detected gRNAs enables greater power than single-plex CRISPRa screens, as any given single-cell transcriptome is informative with respect to multiple perturbations^2^. In this proof-of-concept study, we performed two screens in which the same set of cCREs was targeted, first in K562 cells and then in human iPSC-derived excitatory neurons. We discover both enhancer and promoter-targeting gRNAs capable of mediating upregulation of target gene(s). For enhancers in particular, the upregulatory potential of individual gRNAs was consistently restricted to one cell type, implying a dependency on either the *cis* chromatin landscape and/or additional *trans*-acting factors for successful gene activation.

**Figure 1.**
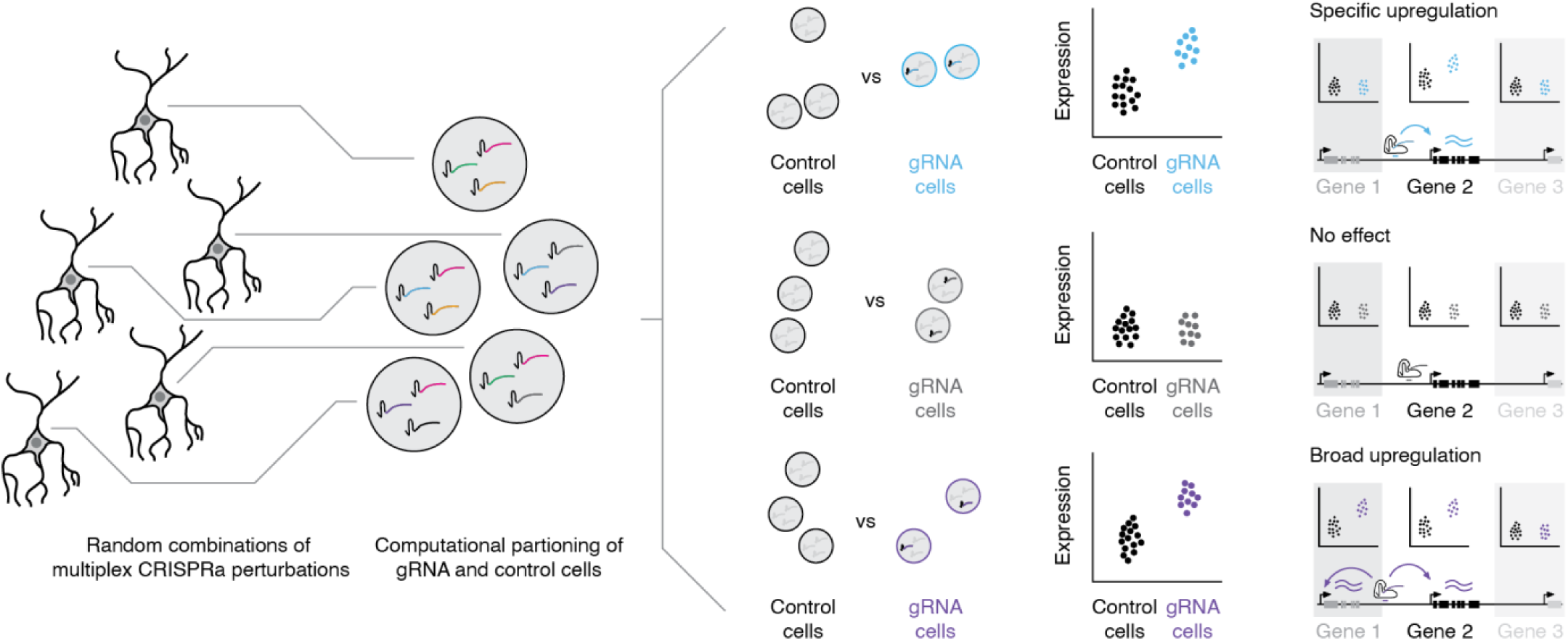
Multiplex, single cell CRISPRa screening for cell type-specific regulatory elements. (Left) A library of gRNAs targeting candidate cis-regulatory elements (cCREs) is introduced in a multiplex fashion to a population of cells expressing CRISPRa machinery, such that each cell contains a random combination of multiple CRISPRa-mediated perturbations. (Middle) Following single cell transcriptional profiling and gRNA assignment, cells are systematically computationally partitioned into those with or without a given gRNA and tested for upregulation of neighboring genes. (Right) CRISPRa perturbations can either result in target-specific upregulation, no detectable effect (*e.g.*, for non-targeting controls) or, at least theoretically, broad *cis*-upregulation of multiple genes in the vicinity of the gRNA/CRISPRa machinery. Furthermore, patterns of upregulation can either be general or cell type-specific.

## Results

### Multiplex single-cell CRISPRa screening of regulatory elements in K562 cells

As a proof of principle, we first sought to implement multiplex single-cell CRISPRa screening in the chronic myelogenous leukemia cell line K562, an ENCODE Tier 1 cell line^15^ in which we had previously performed a multiplex CRISPRi screen^2^. Our proof-of-concept library included gRNAs targeting transcription start site (TSS) positive controls (30 gRNAs), candidate promoters (313 gRNAs), candidate enhancers (100 gRNAs) and non-targeting controls (NTCs; 50 gRNAs). The 30 TSS positive control gRNAs were selected from a previously reported hCRISPRa-v2 library^16^, while the 313 candidate promoter-targeting gRNAs were designed to 50 annotated TSSs of 9 high-confidence haploinsufficient risk genes associated with ASD and NDD (*BCL11A*, *TCF4*, *ANK2*, *CHD8*, *TBR1*, *SCN2A*, *SYNGAP1*, *FOXP1*, and *SHANK3*) that are potential therapeutic targets for CRT^17^. Genes for which haploinsufficient (*i.e.* dominant, loss-of-function) variants appeared to be primary drivers of risk were prioritized. Further prioritization was based on gene cDNA size being too large to fit into traditional gene therapy vectors. The candidate enhancer-targeting guides included 50 gRNAs designed to target 25 enhancer hits previously validated by CRISPRi^2^, as well as 50 gRNAs designed to target 25 enhancer “non-hits” (*i.e.* sequences with biochemical markers strongly predictive of enhancer activity in K562 cells that did not alter gene expression when targeted with CRISPRi^2^) (**Fig. S1a-b**; **Methods**). We cloned this gRNA library (n=493) into piggyFlex, a piggyBac transposon-based gRNA expression vector, to allow for genomic integration and stable expression of gRNAs^18^. The piggyFlex vector has both antibiotic (puromycin) and fluorophore (GFP) markers, enabling flexibly stringent selection for cells with higher numbers of gRNA integrants. We have observed silencing of lentiviral constructs when integrated in iPSCs (data not shown), in part motivating the switch to a transposon-based gRNA delivery vector. Transposon vectors also avoid issues that can arise due to recombination during viral packaging^19^ and associated safety concerns. Additionally, this vector design allows for gRNA transcript capture during single-cell library preparation^18^ (**Fig. S1c**).

There is no consensus on which CRISPRa activation complex is best suited for broad and scalable targeting of enhancers^13^. We therefore tested both the VP64 activation complex, which consists of four copies of the VP16 effector, and the VPR activation complex, which consists of the VP64 effector fused to the p65 and Rta effectors^20,21^. The VPR complex has been shown to lead to higher levels of transcriptional activation than that of the VP64 complex. However, this increased upregulation could achieve higher than therapeutically needed expression levels and being much larger than VP64 could impinge on packaging and delivery of gene therapy vectors such as adeno associated virus (AAV)^21^. We generated a monoclonal, stably VP64-expressing K562 cell line, purchased a polyclonal, stably VPR-expressing K562 cell line (**Fig. 2A**; **Methods**), and validated the capacity of these lines for CRISPRa with a minimal cytomegalovirus (CMV) promoter-tdTomato reporter expression assay^22^ (**Fig. S2**).

**Figure 2.**
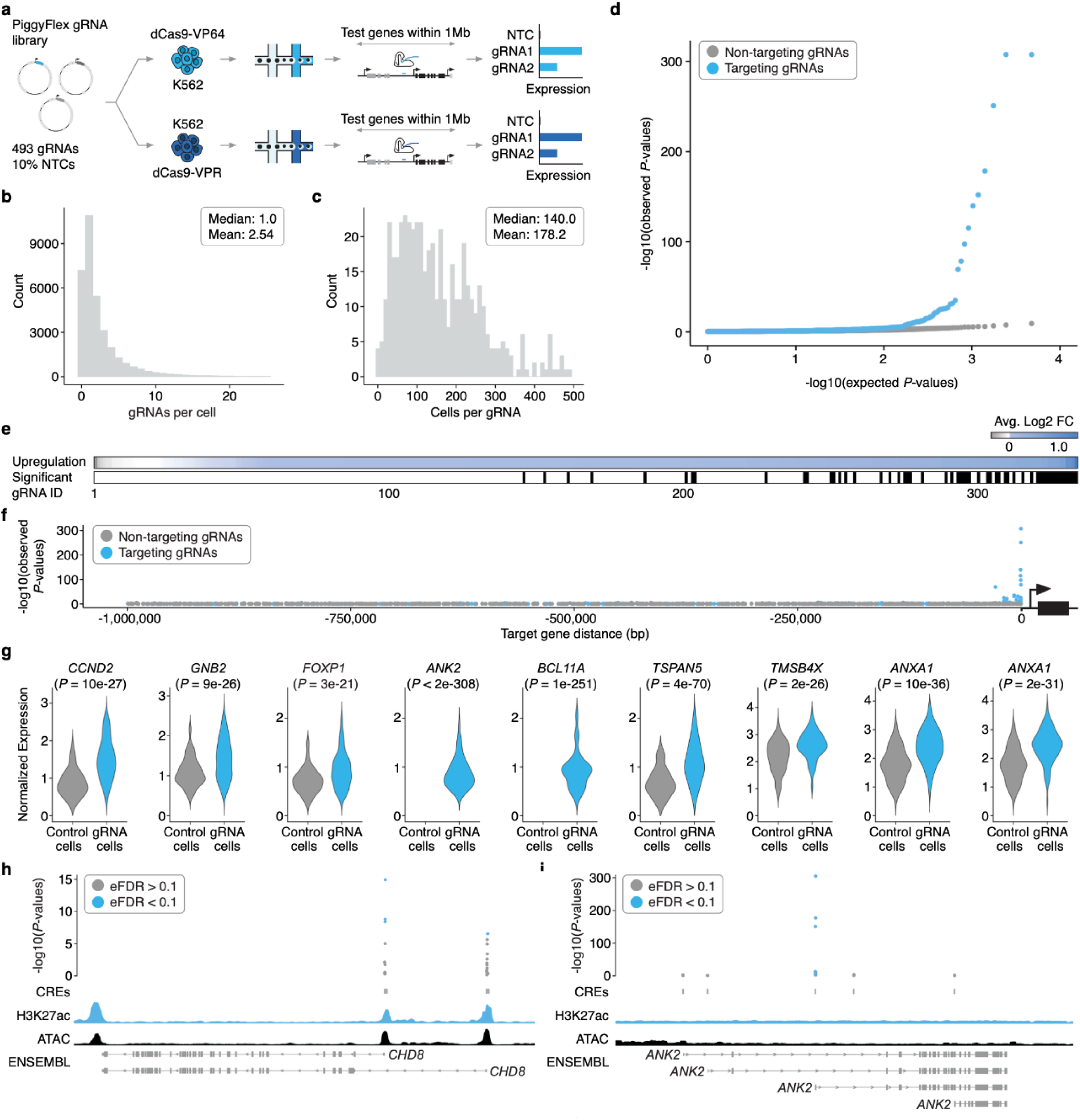
Multiplex single cell CRISPRa screening of regulatory elements in K562 cells. **a)** A piggyFlex library containing gRNAs targeting candidate promoters and distal CREs, TSS positive controls, and 10% NTCs was introduced via nucleofection to two K562 cell lines expressing integrated CRISPRa machinery: 1) K562 CRISPRa-VP64 and 2) K562 CRISPRa-VPR. Following selection, 20,000 cells per CRISPRa K562 line (40,000 total) were harvested and profiled using sc-RNA-seq to capture and assign gRNAs to single cell transcriptomes (see **Fig. S1** and methods for details on piggyFlex design and gRNA capture). **b)** Following QC and gRNA assignment, we identified an average of 2.54 gRNAs/cell (median 1.0 gRNAs/cell). **c)** Multiplexing more than one perturbation per cell enabled an average of 178.2 cells/gRNA (median 140.0 cells/gRNA). **d)** Quantile-quantile plot showing the distribution of expected vs. observed *P*-values for targeting (blue) and non-targeting (gray, downsampled) differential expression tests. **e)** (Top) Heatmap showing the average log2 fold change in expression between cells with each targeting gRNA vs. controls for each of the primary/programmed target genes. Tests are sorted left-to-right by increasing log2 fold change. (Bottom) Categorical heatmap showing which of the perturbations drove significant upregulation using an Empirical FDR approach (EFDR < 0.1). **f)** Targeting gRNAs yielding significant upregulation are enriched for proximity to their target gene. We observe no such enrichment for NTCs tested for associations with target genes randomly selected from the same set. **g)** Example violin plots showing the average log2 fold change between cells with a given gRNA and controls for select hit gRNAs. Hits include TSS positive controls (*CCND2*, *GNB2*), candidate promoters of genes rarely or not expressed in K562 cells (*ANK2*, *BCL11A*) and candidate K562 enhancers (*TSPAN5*, *TMSB4X*, and *ANXA1*). Number of cells bearing each targeting gRNA (from left to right): *CCND2* (n = 73), *GNB2* (n = 220), *FOXP1* (n = 313), *ANK2* (n = 403), *BCL11A* (n = 191), *TSPAN5* (n = 48), *TMSB4X* (n = 260), *ANXA1* (n = 166), *ANXA1* (n = 128). Control cells are downsampled to have the same number of cells as the average number of cells detected per gRNA (n = 178) for visualization. Normalized expression values represent log normalized expression values from Seurat. Only cells with at least 1 target gene UMI are plotted. Note that some genes (*e.g., ANK2*) are typically not detected as expressed in this cell context, resulting in zero UMIs detected and thus no expression distribution plotted in downsampled control cell populations. **h)** Hits included multiple gRNAs targeting isoform-specific promoters of *CHD8*. Empirical *P*-values are visualized alongside tracks for K562 ATAC-seq (ENCODE), H3K27ac signal (ENCODE), and RefSeq validated transcripts (ENSEMBL/NCBI) **i)** The strongest hit gRNAs for ANK2 target the same promoter that is not prioritized by biochemical marks (*e.g.*, accessibility or H3K27ac). Genomic tracks are the same as in panel **h**. Abbreviations: NTC: Non-targeting controls.

We then transfected the gRNA library and piggyBac transposase into each cell line at a 20:1 library-to-transposase ratio to achieve high multiplicity of integration (MOI), and selected cells with puromycin. Cells were cultured for nine days before harvesting for sc-RNA-seq to capture and assign gRNAs to single cell transcriptomes (**Fig. 2a**; **Fig. S1**). After QC filtering, we recovered 33,944 high-quality single-cell transcriptomes across the two cell lines, with 79% of cells having one or more detected gRNAs. We recovered a mean of 2.5 gRNAs per cell (**Fig. 2b**) and 178 cells per gRNA (**Fig. 2c**). Transcriptome quality, MOI, gRNA assignment rate, and gRNA coverage were similar across all four sc-RNA-seq batches (10x Genomics lanes) as well as the two cell lines tested (**Fig. S3**).

To systematically assess the effect of each CRISPR perturbation on target gene expression, we adapted an iterative differential expression testing strategy in which all single cell transcriptomes are computationally partitioned into cells with or without a given gRNA^2^. These two groups are then tested for differential expression of all genes within 1 megabase (Mb) (upper estimate of topologically associated domain size in mammalian genomes^23^) upstream and downstream of the gRNA target site (**Fig. 1**; **Fig. 2a**; **Methods**). In both VP64- and VPR-mediated CRISPRa screening experiments, we observed clear upregulation from both promoter and enhancer-targeting gRNAs (276/391 log2FC>0, 70.6%, *P*<2.2×10^-16^, Fisher’s Exact Test; **Fig. 2d-e**). The presence of an excess of highly significant *P*-values for cells harboring targeting gRNAs versus non-targeting controls (NTCs) also indicates that this multiplex framework successfully detects upregulation of genes from CRISPRa perturbations (**Fig. 2d**). Effects were consistently much stronger and more significant in the dCas9-VP64 cell line as compared to the dCas9-VPR line (**Fig. S3**). This may be due to differences between the VP64 and VPR effectors, site-of-integration effects (VP64 line is monoclonal while VPR line is polyclonal), MOI differences of the integrated effectors, power differences (more cells were recovered per perturbation for the VP64 line than the VPR line), or a combination of these factors.

To identify significant associations between cCRE-targeting gRNAs and their target genes, which we term “hit gRNAs”, we set an empirical false discovery rate (FDR) threshold based on the *P*-values from the NTC gRNA differential expression tests, which are subject to the same sources of noise and error as the targeting gRNA tests. Using an empirical FDR cutoff of 0.1 (**Methods**), we identified 59 activating gRNA hits, including 8 TSS-targeting positive control gRNAs, 39 candidate promoter-targeting gRNAs, 9 distal enhancer hit gRNAs, 2 distal enhancer hit gRNAs wherein the target gene of CRISPRa vs. CRISPRi differed, and 1 distal enhancer non-hit gRNA (in the last three contexts, hit vs. non-hit refers to whether they were “hits” in the previous CRISPRi-based screen with the same guides and cell line^2^) (**Fig. 2e**; **Fig. S4**). Successfully activating gRNAs were strongly enriched for targeting regions proximal to the genes that they upregulated (**Fig. 2f**) and were specific to their predicted target (45/47 promoter-targeting gRNA hits and 9/12 successful enhancer-targeting gRNAs exclusively upregulated the predicted target and no other gene within 1 Mb; **Fig. S4**; **Tables S2-S4**). The gRNAs that upregulated a gene other than the predicted target are discussed further below. Of note, we also observed no instances where targeting a regulatory element, whether a promoter or enhancer, caused significant upregulation of >1 gene.

Taken together, these results demonstrate the potential of this framework to efficiently identify promoter- or enhancer-targeting gRNAs that drive specific upregulation of their target genes in a cell type of interest. Of note, the promoters that were successfully targeted with CRISPRa included genes that were already well-expressed (*e.g., CCND2*, *GNB1*), including two that are haploinsufficient neurodevelopmental disease genes (*FOXP1*, *CHD8*) (**Fig. 2g-h**; **Fig. S4**; **Fig. S5**; **Tables S2-S4**). For *CHD8*, in which variants leading to haploinsufficiency are important risk factors for ASD and NDD^24,25^, we identified multiple CRISPRa gRNAs targeting distinct isoform-specific promoters, providing a potential inroad to isoform-specific CRT (**Fig. 2h**; **Fig. S4**).

Our strongest hits were at the promoters of genes with very low or undetectable expression in K562 (*e.g., ANK2*, *BCL11A*; **Fig. 2g**; **Fig. S5**; **Tables S2-S4**). For example, we identified multiple CRISPRa gRNAs targeting *ANK2*, an ASD/NDD risk gene with a complex isoform structure^24,25^ that is very lowly expressed in K562 cells (**Fig. 2i**). Interestingly, the strongest hits for *ANK2* all targeted a TSS that is not prioritized by biochemical marks (*i.e.*, it is relatively inaccessible and displays a low degree of H3K27ac in K562 cells compared to candidate TSSs of other genes in our library; **Fig. 2i**). On the other hand, for many targeted TSSs or promoters, only one gRNA, if any, activated their target gene when coupled to CRISPRa. More specifically, out of the 313 candidate promoter-targeting gRNAs designed to 50 annotated TSSs of 9 genes, only 37 gRNAs, targeting 11 TSSs and 5 genes, successfully mediated upregulation. One gRNA upregulated a different gene (*WWC3*) than the intended target (*FOXP1*). These results underscore the value of inclusive, empirical screens to identify both CRISPRa-competent promoters as well as gRNAs that can successfully activate them.

At the outset of this work, it was unclear if targeting CRISPRa perturbations to enhancers alone (without co-targeting putatively associated promoters) could reliably increase target gene expression to an extent detectable with conventional sc-RNA-seq^9,12,13^. To determine if CRISPRa targeted to a single enhancer alone could effectively upregulate target gene expression, we analyzed our 50 targeted candidate enhancers, 25 of which were previously validated by multiplex CRISPRi in K562 cells^2^. We observed target gene upregulation for 8 of these 50 targeted candidate enhancers (as noted above, mediated by 12 gRNAs; **Fig. 2g**; **Fig. S4**; **Fig. S5**). Six of the 8 enhancers come from the set of 25 enhancer-gene pairs that we also identified with CRISPRi^2^, including several cases where distinct gRNAs targeting the same enhancer are both successful, *e.g.* two CRISPRa-competent enhancers of *ANXA1* (**Fig. 2g**; **Fig. S4**; **Fig. S5**). In addition, we identified: (1) an enhancer-targeting gRNA that was not a hit in the CRISPRi screen, but here led to upregulation of *HMGA1*; and (2) two enhancer-targeting gRNAs that mediate downregulation of *TUBA1A* when coupled to CRISPRi, but upregulation of *ASIC1* when coupled to CRISPRa.

*ASIC1* is not typically detected as expressed in K562 cells, explaining why e-ASIC1 was not detected as necessary for *ASIC1* expression via CRISPRi. In other words, this enhancer was detected as necessary for high *TUBA1A* expression (an enhancer-gene pair discovered with CRISPRi) but sufficient for increasing *ASIC1* expression when targeted with CRISPRa. While the precise mechanisms underlying this differential behavior of e-ASIC1 in relation to these two genes remains to be characterized, it may be that be that *ASIC1* requires a higher degree of stimulation (in this case, introduced via a CRISPRa activation), whereas *TUBA1A* is activated sufficiently via the same enhancer at a lower level of stimulation (i.e. at the baseline resting state of K562 cells). Alternatively, the *TUBA1A* link may be a false positive of the CRISPRi study^2^, which used an empirical 10% FDR for identifying hits.

Taken together, these results show that multiplex CRISPRa screens leveraging sc-RNA-seq can identify enhancer-targeting gRNAs that can mediate upregulation of specific genes without co-targeting of the corresponding promoters (**Fig. 2g**; **Fig. S4**; **Fig. S5**; **Tables S2-S4**). Furthermore, differences in activity and target-choice despite using the same gRNAs hint at potential differences between CRISPRi and CRISPRa that warrant further exploration.

### Multiplex single-cell CRISPRa screening of regulatory elements in post-mitotic iPSC-derived neurons

We next sought to extend this framework beyond K562 cells to a model that is more relevant for native biology as well as CRT, post-mitotic human induced pluripotent stem cell (iPSC)-derived neurons (**Fig. 3a**)^26^. We initially attempted to generate an iPSC line with CRISPRa-VP64 machinery integrated randomly via lentiviral transduction. However, monoclonal lines generated with this approach silenced the CRISPRa-VP64 machinery during neural differentiation, preventing use in our screening framework. To circumvent this, we used a WTC11 iPSC line equipped with a doxycycline-inducible *NGN2* transgene expressed from the *AAVS1* safe-harbor locus to drive neural differentiation, as well as a ecDHFR-dCas9-VPH construct, expressed from the *CLYBL* safe-harbor locus, to drive CRISPRa (**Fig. S6A-B**)^6^. In this line, addition of doxycycline to induce *NGN2* expression and trimethoprim (TMP) to inhibit the ecDHFR degrons drives neural differentiation and initiates CRISPRa^6^. Expression of NGN2 in iPSCs commits these cells to a neuronal fate, and post-mitotic neurons with neuronal morphology develop within days^27^.

**Figure 3.**
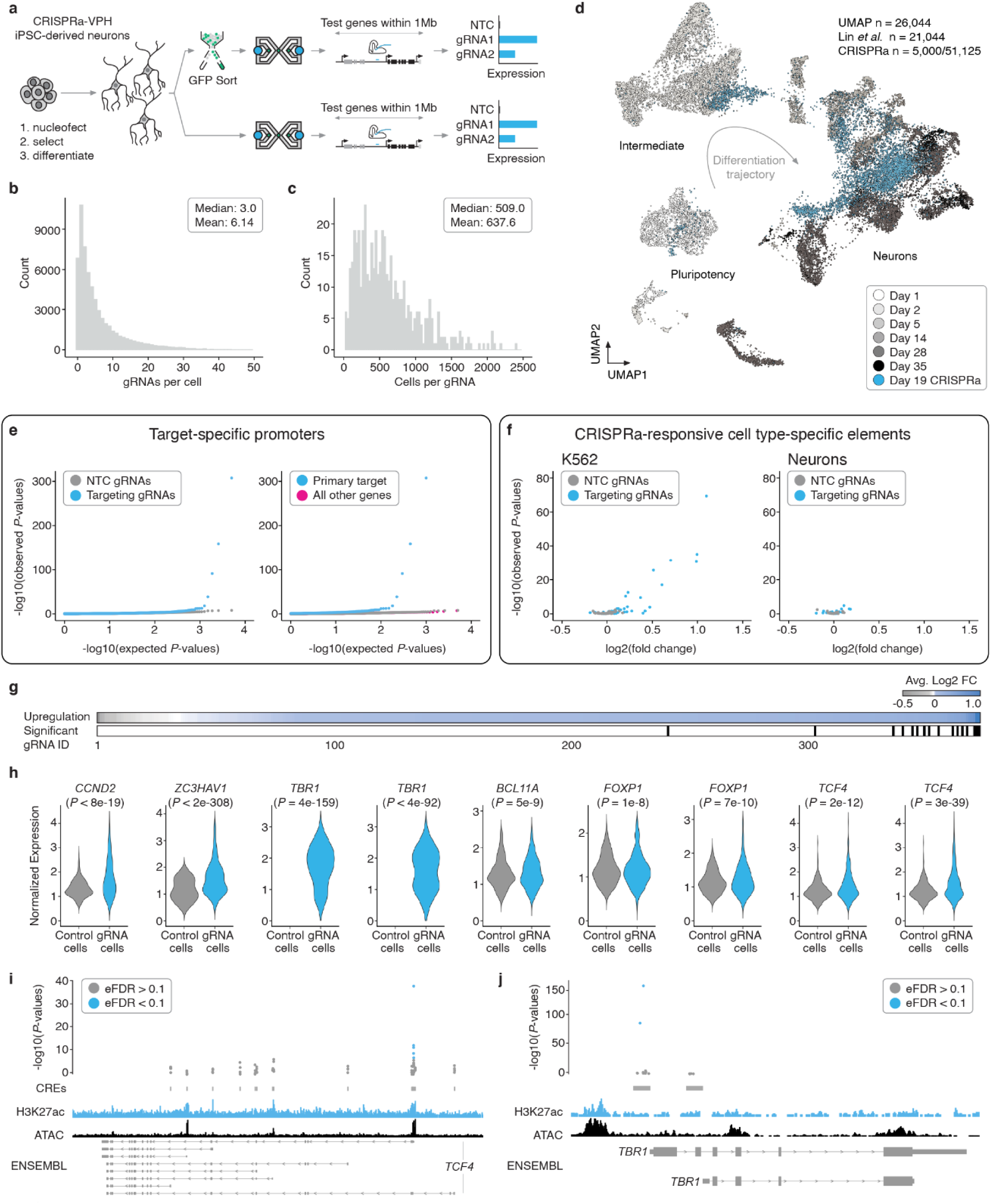
Multiplex single cell CRISPRa screening of regulatory elements in post-mitotic iPSC-derived neurons. **a)** The same piggyFlex library as used in K562 experiments was introduced to a human WTC11 iPSC line harboring TMP-inducible CRISPRa machinery and a Dox-inducible NGN2 transgene to drive neural differentiation. Following selection and differentiation, cells were harvested and profiled with sc-RNA-seq to capture gRNAs and assign them to single cell transcriptomes. Half of the neurons were sorted on GFP immediately prior to sc-RNA-seq to increase the multiplicity of gRNA integrations. **b)** Following QC and gRNA assignment, we identified an average of 6.14 gRNAs/cell (median 3.0). **c)** Neuron gRNA coverage: each gRNA was identified in an average of 637.6 cells (median 509.0). **d)** UMAP projection of the neuron dataset from this study (blue, 51,183 cells downsampled to 5,000 cells to aid with visualization) onto a sc-RNA-seq differentiation time-course from a similar differentiation protocol and NGN2 iPSC line (21,044 cells)^28^. This reference time-course dataset is coloured from white to black based on differentiation day. **e)** (Left) QQ-plot displaying observed vs. expected *P*-value distributions for targeting (blue) and NTC (downsampled) populations. (Right) QQ-plot for targeting tests against their intended/programmed target (blue) compared to targeting tests of all other genes with 1Mb of each gRNA (pink) and NTCs (gray downsampled). There is a clear excess of highly significant *P*-values for programmed targets compared to targeting tests of neighboring genes (pink) or NTCs (gray). **f)** Volcano plot showing the average log2 fold change and *P*-values exclusively for gRNAs that target putative enhancers in K562 cells (left) and iPSC-derived neurons (right). **g)** (Top) Heatmap showing the average log2 fold change in expression between cells with each targeting gRNA vs. controls for each of the primary/programmed target genes. (Bottom) Categorical heatmap showing which of the perturbations produced significant upregulation using an Empirical FDR approach (EFDR < 0.1). **h)** Example violin plots showing the average log2 fold change between cells with a given gRNA and controls for select hit gRNAs. Hits include TSS positive controls (*CCND2*, *ZC3HAV1*), candidate promoters of genes rarely or not expressed NGN2, including the cortical neuron marker *TBR1*, and candidate promoters of genes with native expression in iPSC-derived neurons that could be further upregulated (*BCL11A*, *FOXP1*, and *TCF4*). Number of cells bearing each targeting gRNA (from left to right): *CCND2* (n = 350), *ZC3HAV1* (n = 765), *TBR1* (n = 455), *TBR1* (n = 655), *BCL11A* (n = 964), *FOXP1* (n = 1198), *FOXP1* (n = 917), *TCF4* (n = 1051), *TCF4* (n = 1253). Control cells are downsampled to have the same number of cells as the average number of cells detected per gRNA (n = 638) for visualization. Normalized expression values represent log normalized expression values from Seurat. Only cells with at least 1 target gene UMI are plotted. Note that some genes (*e.g., TBR1*) are typically not detected as expressed in this cell context, resulting in zero UMIs detected and thus no expression distribution plotted in downsampled control cell populations. **i)** Of 14 targeted candidate promoters, five hit gRNAs for TCF4 target the same candidate promoter that aligns with biochemical marks of regulatory activity (ATAC-Seq and H3K27ac). Empirical *P*-values are visualized alongside tracks for iPSC-derived neuron ATAC-seq (accessibility)^45^, and H3K27ac^45^, and RefSeq validated transcripts (ENSEMBL/NCBI). **j)** Hits included multiple gRNAs targeting *TBR1*. Genomic tracks are the same as in panel **i**.

After optimizing iPSC transfection conditions to achieve high numbers of integrated gRNAs per cell via nucleofection, we integrated the same gRNA library (at a 5:1 gRNA-library:transposase ratio) into iPSCs as we did for the K562 screen (**Fig. 3a**). Following integration, we confirmed functional CRISPRa activity in these neurons via the same tdTomato expression assay used in our K562 CRISPRa validation (**Fig. S6b**). In addition to optimizing transfection conditions, we sought to further boost the multiplicity of gRNA integrations per cell by selecting the cells with a high concentration of puromycin (**Fig. 3a**). After differentiating to neurons over 19 days, we proceeded to sc-RNA-seq. Half of the neurons went directly into sc-RNA-seq (10x Genomics), while the other half were dissociated and flow sorted based on GFP expression (top 40%) prior to sc-RNA-seq, again with the goal of boosting the multiplicity of gRNA integrations (**Fig. 3a**). After quality control filtering, we retained 51,183 single-cell transcriptomes, of which we recovered 1+ associated gRNAs for 89%. With our optimized transfection protocol, we identified a mean of 6.1 gRNAs/cell (**Fig. 3b**) and a mean of 638 cells that harbored each individual gRNA (**Fig. 3c**). Sorting on GFP expression prior to sc-RNA-seq resulted in a 2-fold increase in the number of gRNAs identified per cell (**Fig. S7**).

Our differentiated neurons most closely resemble 14- to 35-day differentiated neurons obtained via *NGN2* induction in iPSCs by an independent group^28^ (inferred by integration of these sc-RNA-seq datasets; **Fig. 3d**; **Fig. S8**). A minority of the neurons transcriptionally resemble an intermediate neuronal fate, a difference that we tentatively attribute to the absence of co-cultured glia in our differentiation protocol. Although glia are known to promote maturation of NGN2-induced neurons (and were used in generating the dataset we are comparing to^28^), we excluded them because they can also introduce culture variability due to batch effects introduced by primary glia^27^.

We confirmed that the neurons had progressed beyond a pluripotent state and were committed to a post-mitotic neuronal fate by the expression of the pan-neuronal marker *MAP2* and the lack of expression of the pluripotency marker *NANOG* (**Fig. S8**). These neurons also express *LHX9* and *GPM6A*, markers of central nervous system (CNS) neurons (**Fig. S8c**), and *CUX1* and *SLC17A7*, but not GABAergic markers *GAD1* and *GAD2,* supporting their assignment as excitatory rather than inhibitory neurons (**Fig. S8f**)^26^. Consistent with this, when we co-embedded our transcriptome data onto data from Lin et al.^28^, they overlay with “Fate 2” and “Fate 3” cells, which transcriptionally resemble CNS neurons (**Fig. 3d**; **Methods**). Of note, there was no readily apparent enrichment of specific gRNAs within particular clusters (**Fig. S9**), which is consistent with the specificity and modest fold changes of the observed instances of upregulation (**Fig. S9**).

We applied the same differential expression testing strategy as used for the K562 screen to the iPSC-derived neuron screen data, with an empirical FDR cutoff of 0.1 to call significant hits. Similarly to the K562 screen, we observed clear upregulation from targeting gRNAs (281/383 log2FC>0, 73.4%, *P*<2.2×10^-16^, Fisher’s Exact Test) and an excess of highly significant *P*-values for targeting gRNA tests compared to NTCs (**Fig. 3e**), confirming that this overall framework is transferable to more physiologically and clinically relevant models such as iPSC-derived neurons. As with the K562 screen, we observed strong enrichment for genomic proximity between successful gRNAs and their target genes, but no such enrichment for NTCs tested for associations with target genes randomly selected from the same set (**Fig. S10**).

There were 17 hit gRNAs in neurons (FDR < 0.1; **Fig. 3g**), all of which were TSS-targeting positive controls (n = 6) or candidate promoters of ASD/NDD risk genes (n = 11) (**Fig. S11**). Of these 17 hit gRNAs, 12 were also hits in the K562 screen while 5 were specific to iPSC-derived neurons (**Fig. S12a**). The screen in iPSC-derived neurons was strikingly target-specific: 16 of 17 of our identified hits, all promoter-targeting gRNAs, upregulated their anticipated target gene and no other genes within the 1-Mb window tested (**Tables S5-S7**). The only gRNA hit in iPSC-derived neurons resulting in upregulation of an unintended gene was a gRNA targeting the TSS of the pseudogene *PPP5D1* that led to upregulation of the calmodulin gene *CALM3* (**Fig. S11d**), but this is presumably due to these two genes sharing a bidirectional, outward-oriented core promoter. This gRNA also drove upregulation of *CALM3* in the CRISPRa screen of K562 cells (**Fig. S4d**). These results, in combination with the K562 results discussed earlier, reveal that certain gRNAs designated ‘TSS-targeting positive controls’ based on their ability to yield a growth phenotype in bulk screens of cancer cell lines, do not yield strong upregulation of their putative target gene (**Fig. S4c; Fig. S11c**). We observed no significant differences across several characteristics (*e.g.*, GC content, baseline target gene expression level, the number of cells harboring each gRNA) between gRNAs yielding successful activation and those not in K562 cells and neurons, with the exception that K562 enhancer hit gRNAs tended to have more cells (**Fig. S13**). However, upon conducting an analysis of epigenetic features available for both cell lines as well as for *in vivo* developing brain samples, we found that promoter- and enhancer-targeting hit gRNAs were more likely to fall within regions of active, open chromatin and were conversely depleted from regions with repressive marks (**Fig. S14**; **Table S8**).

Similar to K562 cells, we observed several instances where a specific TSS was most amenable to activation (**Fig. S15**). One such example is *TCF4*, an ASD/NDD risk gene^24,25^that is a strong candidate for CRT due to its large cDNA size (precluding it from fitting into an AAV) and complex locus architecture. We tested 14 candidate TSSs of *TCF4* and identified 5 gRNAs capable of driving upregulation of *TCF4* in neurons, all of which target the same candidate TSS that resides in open chromatin with strong H3K27ac signal (**Fig. 3h-i**; **Fig. S15a**; **Fig. S16**). Our hits also included examples of cell type-specific promoters. Among these were several gRNAs targeting candidate promoters of ASD/NDD risk genes capable of upregulating genes that are not expressed or rarely expressed in iPSC-derived *NGN2*-differentiated neurons (**Fig. 3h**; **Fig. S16**). For example, gRNAs targeting the promoter of *TBR1*, a transcription factor expressed in forebrain cortical neurons but known not to be expressed in NGN2-differentiated iPSC-derived neurons^26^ led to *TBR1* upregulation (**Fig. 3j**; **Fig. S15b**; **Fig. S16**). Of note, these same gRNAs did not result in upregulation of *TBR1* in K562 cells. This suggests that these neurons are in a permissive state for CRISPRa to activate *TBR1*, despite a lack of highly accessible chromatin in the region targeted by the *TBR1* gRNA (**Fig. 3h,j**; **Fig. S15b**). Whether these differences in “*TBR1* activatability” are due to differences in the chromatin environment at this locus between K562 cells and iPSC-derived neurons, or alternatively differences in the milieu of *trans*-acting factors, remains an open question.

However, in contrast to the cell type-specific promoter examples noted above, we more often observed consistent upregulation across promoter targets and TSS-targeting controls between the two cell types (**Fig. S12**). Specifically, 12 out of 17 of the promoter- and TSS-targeting hit gRNAs in neurons were also hits in K562 cells, and upregulation was correlated across cellular contexts (Pearson’s correlation coefficient = 0.75, **Fig. S12**). In contrast, we observed striking cell type-specificity for targeted enhancers that were successfully upregulated. While 20% (12/60) of our K562 screening hits were enhancer-targeting gRNAs (**Fig. S4**), none of these were also hits in neurons (**Fig. S11**; **Fig. S17**). Even putting aside significance, the fold-effects on the anticipated target genes of K562-competent activating gRNAs were not well-correlated between cell types (**Fig. 3f**; **Fig. S12b,** Pearson’s correlation coefficient = −0.18). Overall, these results show that it is possible to drive cell type-specific upregulation of a gene of interest by targeting CRISPRa to a cell type-specific distal enhancer, without co-targeting of the corresponding promoter.

### Reanalysis with covariates and singleton experimental validations support and extend results

To better account for covariates, we applied SCEPTRE^29–32^, an independently developed analytical framework based on conditional resampling that integrates various covariates to calibrate the statistical assessment of results in single cell CRISPR screens (**Methods**). SCEPTRE detected clear signals in both datasets (**Fig. S18**; **Fig. S19**; **Tables S9-14**). Encouragingly, SCEPTRE results aligned well with our original approach, but yielded more hits. Our original approach identified 59 activating hit gRNAs in K562 cells and 17 activating hit gRNAs in neurons, while SCEPTRE found 83 K562 hits and 35 neuron hits. 48/59 K562 (81%) and 15/17 neuron (88%) hits found with our approach were also found by SCEPTRE (**Fig. S18**; **Fig. S19**; **Tables S9-14**). The few cases where our original approach found a hit that SCEPTRE did not, were primarily due to the fact SCEPTRE has a more stringent minimum threshold on the proportion of cells required to show expression of the target gene (*e.g., ASIC1* expression is rarely detected in K562 cells, yet two independent gRNAs targeting a distal enhancer led to clear upregulation in our screen; see also singleton validations below).

Several notable findings arose from this analysis. First, SCEPTRE identified a single hit gRNA yielding upregulation of *SCN2A* exclusively in iPSC-derived neurons out of 20 *SCN2A*-targeting gRNAs tested (**Fig. S17**; **Table S14**). This is the same “H1 gRNA” we recently used in therapeutic development rescue studies in *SCN2A+/-*haploinsufficient mice and human iPSC-derived neurons^11^, providing clear evidence our scalable single cell framework can identify therapeutically relevant gRNAs. Second, despite more than double the hits in neurons (35 neuron hits with SCEPTRE vs. 17 with our original approach), and several new enhancer hits in K562 cells (21 enhancer hits with SCEPTRE vs. 12 with our original approach), the signal from K562 enhancers is still overwhelmingly cell type-specific (SCEPTRE calls only 1 enhancer hit in neurons vs. 21 enhancer hits in K562 cells) (**Fig. S18**; **Fig. S19**; **Tables S9-14**).

To further validate our results, we selected 8 of our hit gRNAs to test in singleton experiments. Validation gRNAs were selected to represent a range of significance levels, as well as both shared and cell-type specific promoters and enhancers (**Table S15**). We cloned the 8 gRNAs as singleton constructs, introduced them at high MOI to create 3 independent polyclonal cell lines in each cell context, selected, and in the case of neurons, differentiated cells, and then created bulk RNA-seq libraries for each line (8 gRNAs x 2 cell contexts x 3 replicate cell lines = 48 independent polyclonal lines and corresponding bulk RNA-seq samples). We then compared expression levels in the 3 replicate lines with a given targeting gRNA to expression levels in the remaining 21 lines from that cell context as a control (mirroring our single-cell analysis framework).

We observed consistent, clear upregulation of our intended target genes across both cell contexts in nearly all cases. Specifically, 14/16 validation experiments (7/8 in each cell context, 88%) showed the expected upregulation, or not, in the corresponding cell type (**Fig. 4**; **Fig. S20**). Further, gRNAs targeting a cell-type specific promoter of *TBR1* drove the expected upregulation exclusively in iPSC-derived neurons (**Fig. 4**). Hit gRNAs targeting e-ANXA1 and e-ASIC1, two enhancers identified as K562-specific in our single-cell analysis, drove upregulation of *ANXA1* and *ASIC1* exclusively in K562 cells in singleton validations (**Fig. 4**). Of note, e-HMGA1, a putative shared enhancer, drove the expected upregulation in neurons but not in K562 in singleton validations, so we removed it from our list of high-confidence, singleton validated CRISPRa-responsive CREs (**Fig. 4**). These experiments were designed to validate upregulations of target genes in targeted tests, though in some cases we were able to detect additional biologically coherent gene expression changes on a genome-wide basis, potentially secondary effects of upregulating the primary target (**Figs. S21, S22; Table S16**). For example, in both K562 cells and iPSC-derived neurons, genes that were significantly downregulated upon CRISPRa of *BCL11A* are highly enriched for genes involved in oxidative phosphorylation (Gene Set Enrichment Analysis (GSEA); FDR q-value < 1e-20). Importantly, fold changes in target gene expression observed in singleton bulk RNA-seq validation experiments correlated well with results from our multiplex single-cell framework (Pearson’s correlation coefficient = 0.85) (**Fig. S23**). Taken together, these results refine and extend our framework and confirm our ability to identify therapeutically relevant, target- and cell type-specific CRISPRa-responsive regulatory elements and gRNAs that target them.

**Figure 4.**
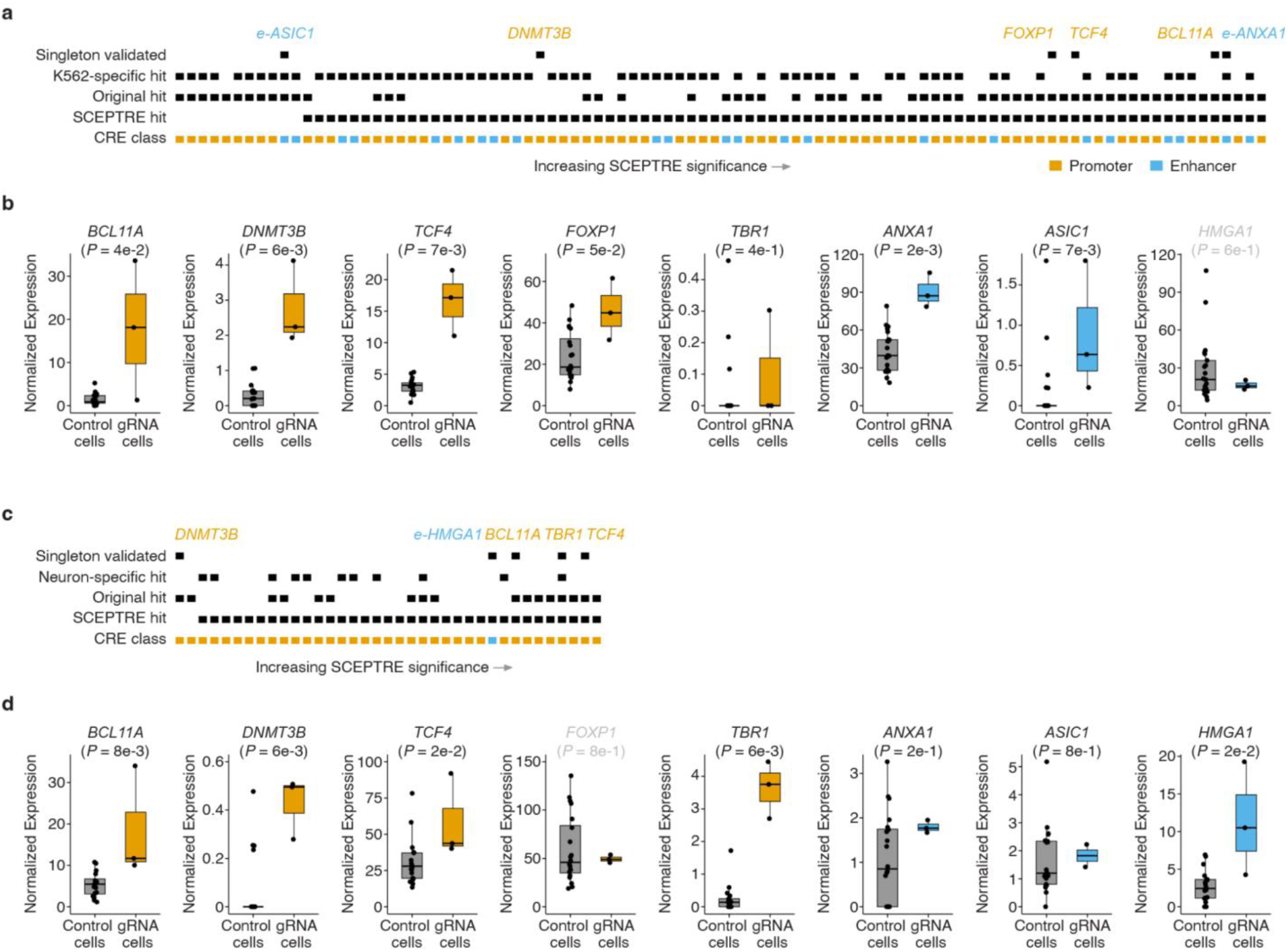
Reanalysis with covariates and singleton experimental validations support and extend results. **a)** Singleton validation results for K562 cells. Categorical heatmap indicating whether a K562 hit gRNA was detected with SCEPTRE, our original approach, targeted a promoter or enhancer, drove K562-specific upregulation, and whether it was validated with singleton experiments. Hit CREs that drove upregulation in singleton validations are labeled and coloured according to target CRE class. **b)** Boxplots displaying the average log2 fold change between cells with a given gRNA (n = 3 independent lines) and controls (n = 21 lines). Normalized expression displayed in transcripts per million (TPM). **c)** Singleton validation results for iPSC-derived neurons. Categorical heatmap indicating whether a neuron hit gRNA was detected with SCEPTRE, our original approach, targeted a promoter or enhancer, drove iPSC-derived neuron-specific upregulation, and whether it was validated with singleton experiments. Hit CREs that drove upregulation in singleton validations are labeled and coloured according to target CRE class. **d)** Boxplots displaying the average log2 fold change between cells with a given gRNA (n = 3 independent lines) and controls (n = 21 lines). Normalized expression displayed in transcripts per million (TPM). *P*-values are from a two-tailed Wilcoxon rank-sum test with a significance threshold of 0.1. The two instances where expected upregulation was not observed in a particular cell context are labeled in gray above the corresponding box plot (specifically gRNAs targeting e-HMGA1 in K562 cells and a promoter of *FOXP1* in neurons). The 14/16 experiments where the expected result was observed are labeled in black above the corresponding box plot, while the 2/16 that are labeled in gray failed to validate.

## Discussion

Here we describe a scalable, multiplex single-cell CRISPRa screening framework to identify cell type-specific CREs, the genes they regulate, and gRNAs that successfully target them for use in CRT. In applying this framework, we identified gRNAs functionally and cell type-specifically targeting endogenous CREs of haploinsufficient genes in K562 cells and iPSC-derived excitatory neurons. To our knowledge, this is the first screen of this scale and kind (highly multiplexed, single-cell resolution noncoding CRISPRa perturbations) performed successfully in a post-mitotic, therapeutically relevant cell type. Our results include several examples of enhancer-target gene relationships that are experimentally supported in their native genomic context, which remain exceedingly rare in the literature due to the significant technical barriers that have persisted despite major efforts to discover or confirm more such relationships. Importantly, our approach lays the groundwork for discovering hundreds to thousands more endogenous enhancer-gene relationships in non-workhorse cancer cell lines such as post-mitotic iPSC-derived neurons. For example, we anticipate it will be possible to massively scale screening for gRNAs and cell-type-specific CREs capable of upregulating remaining functional copies of the roughly 660 genes known to cause disease when haploinsufficient.

In principle, increases in target gene expression of less than 100% to a few fold, especially if cell-type specific, would be ideal for therapeutic purposes in haploinsufficient disorders^33–35^. Many of our fold-changes meet these criteria (median fold-change for all hits = 1.7; 79/118 SCEPTRE hits achieved at least 50% upregulation, 42/118 hits achieved at least a 2-fold upregulation; 49/118 hits achieved between 1.5-2.5-fold upregulation). Furthermore, even in cases where upregulation is lower or higher than theoretically ideal at the mRNA level, there can still be therapeutic benefit (*e.g.,* in cases where partial rescue of expression is sufficient for phenotypic improvement, or where post-transcriptional mechanisms buffer overcorrection of expression levels)^33,34^.

Several of our strongest gRNA hits were not prioritized by typical predictors of enhancer function, such as chromatin accessibility or H3K27ac histone modifications. For example, we are able to upregulate *TBR1* in iPSC-derived neurons by targeting a promoter region that is largely within closed chromatin in this cellular context. Indeed, while measures of proximity, accessibility, and enhancer-related biochemical marks are all strong predictors, none are conclusive or deterministic predictors of regulatory sequence function, either alone or in combination. This underscores the importance of empirical, systematic screens for CRISPRa-responsive regulatory sequences with approaches such as the one described here. Ultimately, a variety of factors including chromatin accessibility and epigenetic modifications, gRNA design and target-specific nuances around CRISPRa-responsiveness, appear to collectively shape the likelihood and extent of success of a given CRISPRa perturbation in a given cellular context. Our framework has now provided a set of validated CRISPRa sgRNAs that could be used as positive controls moving forward in other CRISPRa screens (in at least K562 and iPSC-derived postmitotic neurons; the set that overlap may be the best bet for potential ‘universal’ controls), thereby filling a critical gap. Future scaling of this technology and its application to additional, clinically relevant cell types, may provide rich training sets that may enable derivation of rational CRISPRa gRNA design rules for distal, cell-type-specific gene activation, which, in contrast to promoters and CRISPRi^16,36,37^, are quite lacking at present. Further, these results illustrate the unique potential of noncoding CRISPRa screens to identify regulatory elements that can mediate upregulation of target genes, regardless of whether or not the gene is natively expressed in the cell type of interest or not.

A major question that we sought to answer through these experiments was whether one can target candidate enhancer sequences with a CRISPRa perturbation and observe upregulation of an intended target gene via scRNA-seq. There have been relatively few efforts to apply CRISPRa to enhancers to date, and most have focused on a handful of enhancer regions and typically only measuring changes in expression of a specific nearby gene of interest as a readout^9,12,13^. Recent literature suggests that co-targeting a promoter and the candidate enhancer in question can make the enhancer CRISPRa perturbations more efficient and reliable^13^. Although feasible, co-targeting an enhancer and promoter is less likely to yield cell-type-specific upregulation of target genes -- a likely requirement for effective CRT. Delivery of multiple gRNAs also complicates therapeutic delivery and increases the chances of effects on off-target genes (not to mention off-target cell types). Despite using gRNAs that were optimized for CRISPRi screening in our CRISPRa screen, we observed target gene upregulation for 8 of 25 enhancers that we targeted (32%), showing that one can reliably increase target gene expression by targeting enhancers alone. We imagine that this success rate can be improved via a combination of brute force, *i.e.* testing more gRNAs, improved CRISPRa activation domains^38^, and better CRISPRa-specific gRNA design.

Multiplex single-cell CRISPRa screening is a scalable approach to identifying functional CRISPRa gRNAs that can upregulate intended target genes in either a general or cell-type-specific manner. We introduced multiple perturbations per cell, which increased the power of our assay (*i.e.* a mean of 1 gRNA per cell would have required sc-RNA-seq of >400,000 cells to achieve the same power). For additional context, the increased hands-on time and reagents required to generate, select, and differentiate the 48 cell lines used to validate the 16 gRNAs across the two cell contexts made the costs of the singleton and pooled experiments comparable, despite singleton experiments assessing <5% as many gRNAs. Alternative bulk approaches such as those based on integrated fluorescent reporters or FISH scale linearly with a new set of reagents required for each new candidate target gene.

One potential concern is that multiplex, single-cell CRISPR screens that treat cells as pseudo-replicates have the potential to suffer from an overestimation of differentially expressed genes due to the statistical dependency between cells originating from the same starting population^39^. Our singleton validation results allay this concern, but we cannot fully rule out overestimation. Further, given the ease of generating large numbers of differentiated neurons with *in vitro* human neural cultures, and then sorting on the GFP-positive gRNA expression vector prior to single-cell transcriptome profiling, our approach offers a straightforward way to further boost the number of gRNAs captured per cell. In addition, improvements in methods to capture specific transcripts^40–42^, or to perform scRNA-seq more cheaply^43,44^, may enable considerably larger screens for a given cost.

CRT is a promising, next-generation therapeutic approach that harnesses endogenous gene regulatory circuits to treat genetic disorders^10,11,17^. However, CRT requires an intricate knowledge of the regulatory elements capable of driving target gene upregulation at physiologically relevant levels specifically in affected tissues. We envision that the framework described here can be deployed in increasingly sophisticated *in vitro* and *in vivo* models of human development to discover reagents capable of treating the hundreds of disorders associated with low gene dosage.

## Supplementary figures

**Figure S1.**
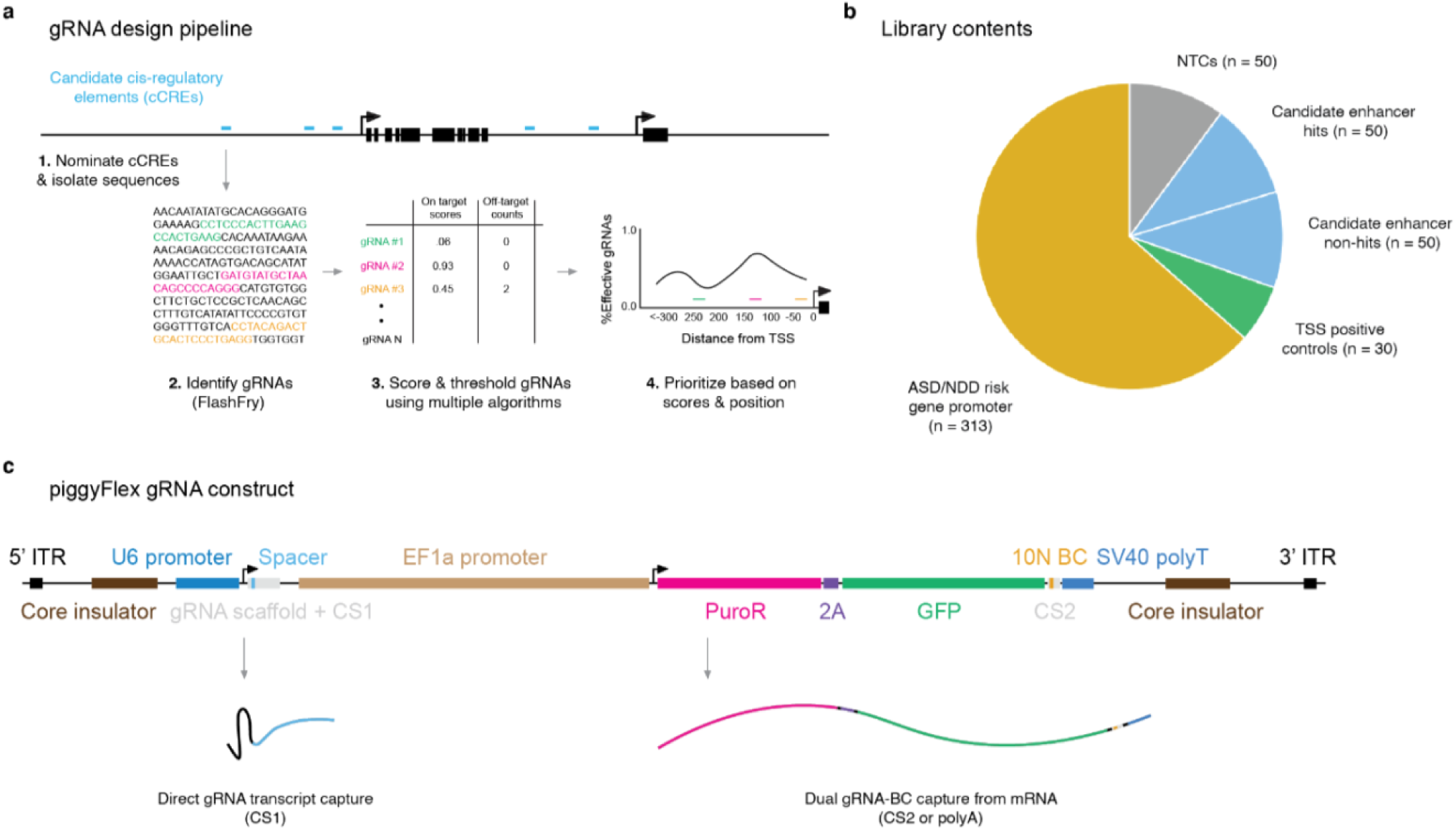
gRNA design pipeline, library contents, and piggyFlex gRNA delivery construct. **a)** gRNA design pipeline. First, candidate Cis-Regulatory Elements (cCREs) surrounding a gene of interest were identified based on biochemical marks of regulatory activity (*e.g.*, accessibility, active transcription, etc.). Next, candidate gRNAs targeting each cCRE were generated using FlashFry^46^. Then, gRNAs were scored and prioritized using multiple algorithms. Finally, in the case of promoters where systematic CRISPRa design rules are available, gRNAs were prioritized based on optimal position relative to the TSS^36^. **b)** PiggyFlex gRNA library contents by target category. **c)** PiggyFlex construct design. PiggyFlex is a piggyBac transposon-based gRNA delivery vector equipped with a dual antibiotic (puromycin) and fluorophore (GFP) selection cassette that enables enrichment for cells with many integrated gRNAs^18^. PiggyFlex enables direct capture of gRNA transcripts or optional capture of gRNA-associated barcodes from GFP mRNA via CS2 or polydT capture.

**Figure S2.**
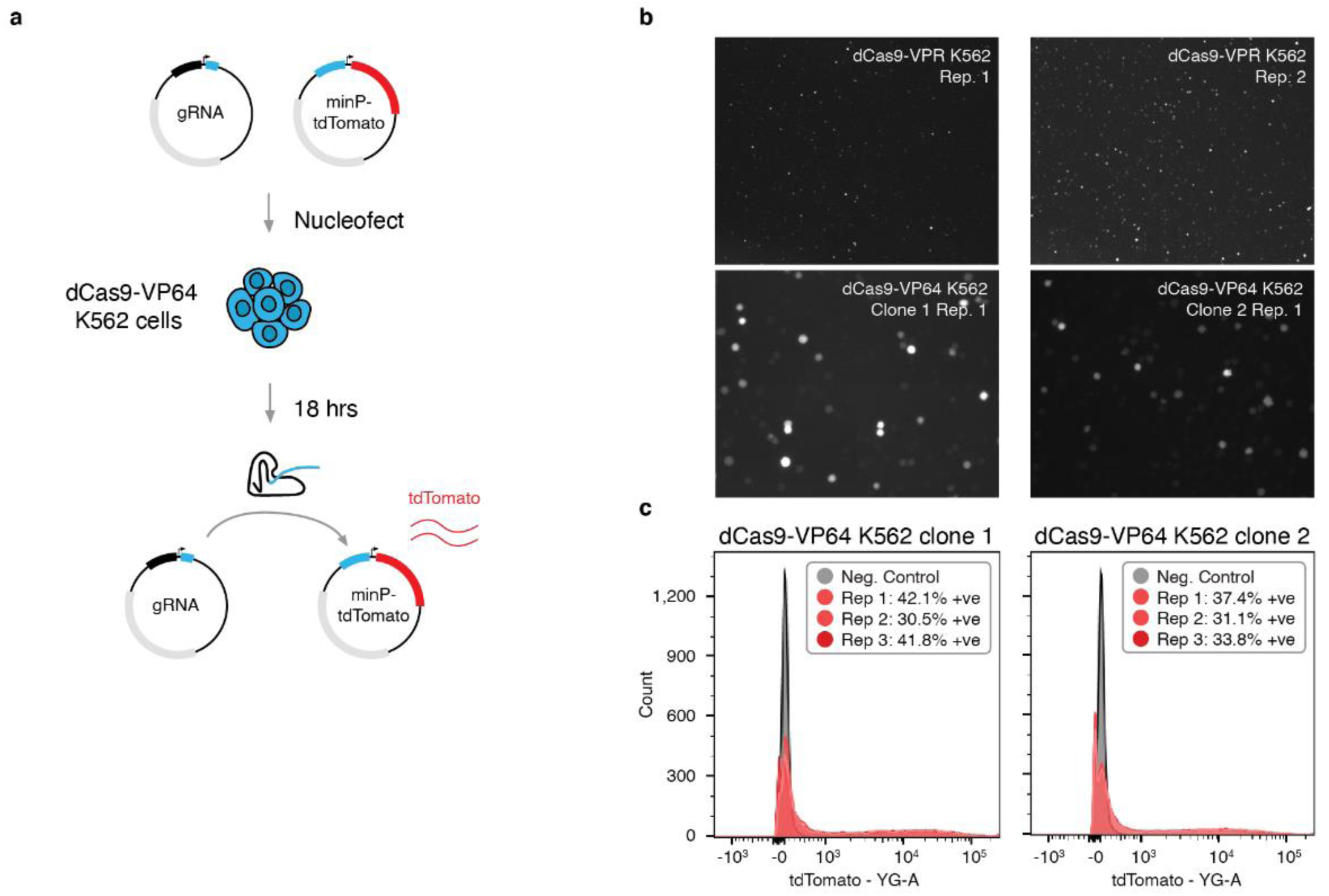
Functional validation of CRISPRa K562 cell lines. a) Schematic of the minP-tdTomato functional assay used to validate CRISPRa cell lines. Two plasmids, one encoding a minP-tdTomato and another encoding a gRNA targeting a sequence immediately upstream of minP were co-nucleofected into K562 cell lines with either dCas9-VP64 or dCas9-VPR constructs integrated (only dCas9-VP64 is illustrated for simplicity). b) Following nucleofection, both dCas9-VPR (top, lower magnification) and dCas9-VP64 (bottom, higher magnification) K562 cell lines drove strong tdTomato expression, confirming the presence of functional CRISPRa machinery in these cell lines. dCas9-VPR images represent two replicate transfections into a single polyclonal line, while dCas9-VP64 images each represent one transfection replicate from two monoclonal lines. Note these are transient transfections without selection, so not all cells are expected to have been successfully transfected and fluoresced under these conditions. c) Example FACS analysis of tdTomato fluorescence in individual dCas9-VP64 transfection replicates of two monoclonal lines.

**Figure S3.**
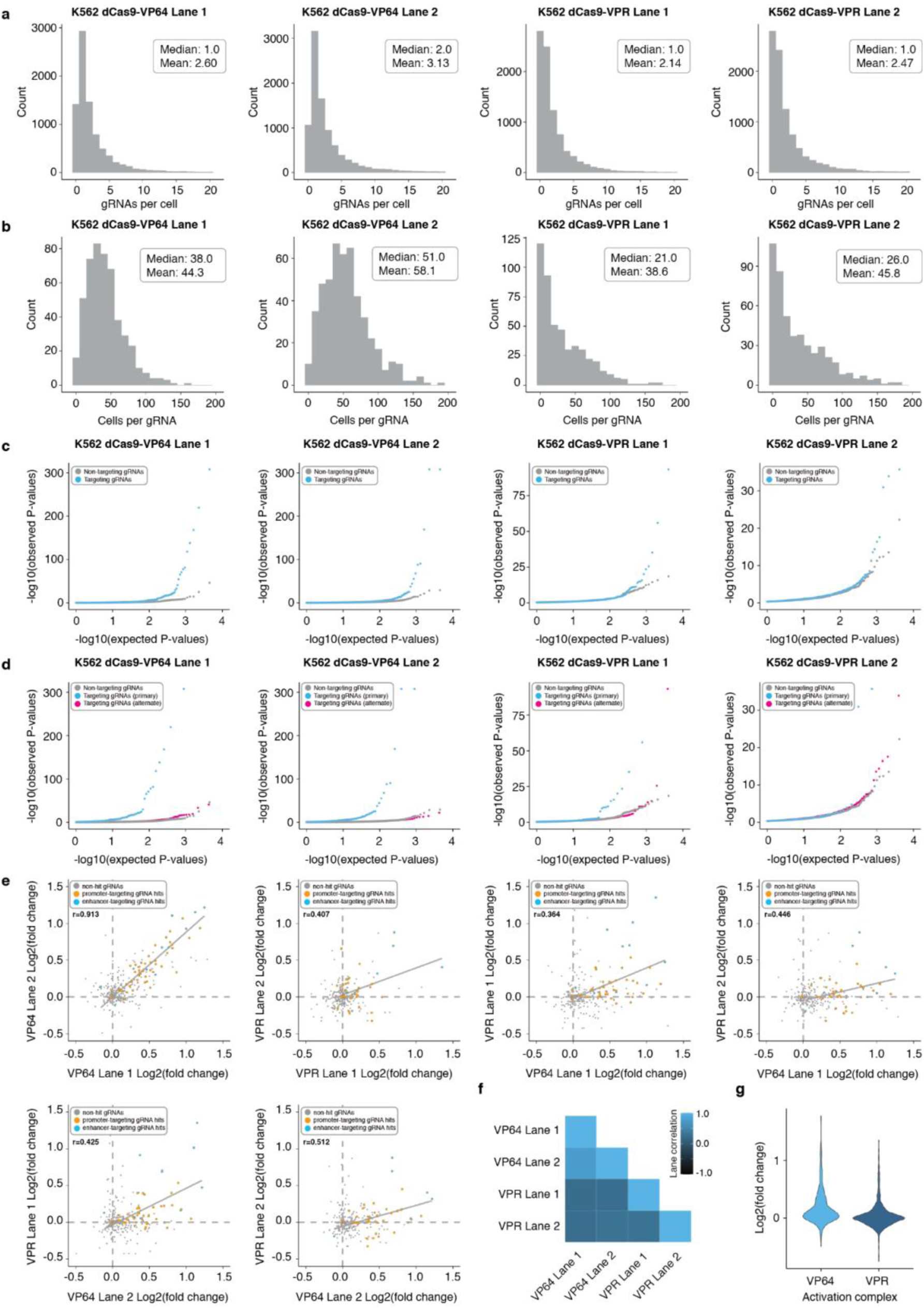
Results for four independent 10x Genomics lanes from K562 screen. a) The four 10x Genomics lanes profiled included two lanes with dCas9-VP64 K562 cells and two lanes with dCas9-VPR K562 cells. Following QC and gRNA assignment we identified an average of 2.60, 3.13, 2.14, and 2.47 gRNAs/cell for the four different 10x Genomics lanes profiled (median 2.60, 3.13, 2.14, and 2.47 gRNAs per cell). PiggyBac integrations per cell distribution is not well-modeled by a standard Poisson distribution and is better approximated by an exponential function. b) Multiplexing more than one perturbation per cell yielded an average of 38.0, 51.0, 21.0, and 26.0 cells/gRNA for the four different 10x Genomics lanes profiled (median 44.3, 58.1, 38.6, and 45.8 cells/gRNA). c) QQ-plots displaying observed vs. expected *P*-value distributions for targeting (blue) and NTC (downsampled) populations across the four different 10x Geomics lanes profiled. d) QQ-plots for targeting tests against their intended/programmed target (blue) compared to targeting tests of all other genes with 1Mb of each gRNA (pink) and NTCs (gray downsampled) across the four different 10x Genomics lanes profiled. e) Correlation plots of log2 fold-changes of gRNAs across the two K562 cell lines (dCas9-VP64 and dCas9-VPR) for all four 10x Genomics lanes profiled. Pearson correlations of gRNA hits are shown. f) Matrix correlation plot displaying the Pearson correlations of the log2 fold-changes of target gene expression values for programmed targets across the four different 10x Genomics lanes profiled. g) Violin plot displaying the log2 fold-changes of target gene expression values for programmed targets for K562 cells harboring the dCas9-VP64 activation complex and the dCas9-VPR activation complex.

**Figure S4.**
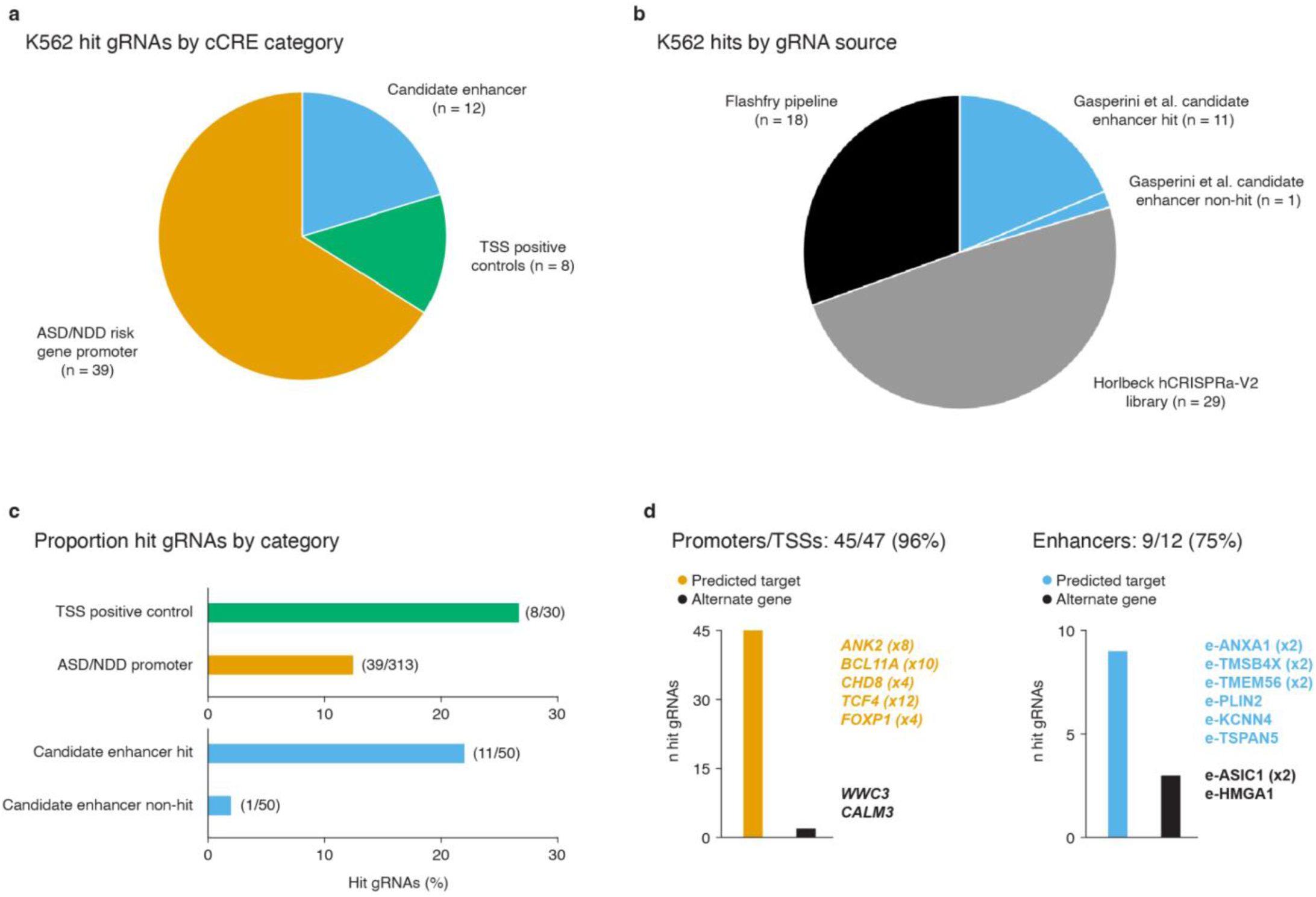
Hit breakdown for screen conducted in K562 cells. a) K562 hit gRNAs by cCRE category. b) K562 hit gRNAs by gRNA source library or design pipeline. c) Proportion of hit gRNAs by cCRE category. d) Proportion of hit gRNAs yielding upregulation of their intended/expected target gene or an alternate gene for candidate promoters/TSSs (left) or enhancers (right). Example hits targeting candidate NDD risk gene promoters (left) and K562 enhancers (right) are listed. Bracketed numbers denote the number of independent hit gRNAs targeting the same cCRE.

**Figure S5.**
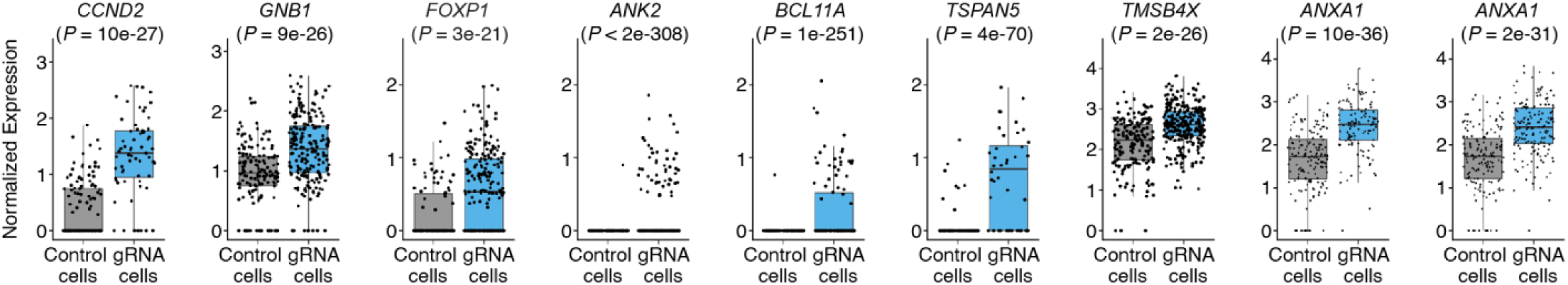
Example upregulations in K562 cells. Example box plots showing the average log2 fold-changes between cells with a given gRNA and controls for select hit gRNAs. Hits include TSS positive controls (*CCND2*, *GNB2*), candidate promoters of genes rarely or not expressed in K562 cells (*ANK2*, *BCL11A*) and candidate K562 enhancers (*TSPAN5*, *TMSB4X*, and *ANXA1*). Number of cells bearing each targeting gRNA (from left to right): *CCND2* (n = 73), *GNB2* (n = 220), *FOXP1* (n = 313), *ANK2* (n = 403), *BCL11A* (n = 191), *TSPAN5* (n = 48), *TMSB4X* (n = 260), *ANXA1* (n = 166), *ANXA1* (n = 128). Control cells are downsampled to have the same number of cells as the average number of cells detected per gRNA (n = 178) for visualization. Normalized expression values represent log normalized expression values from Seurat.

**Figure S6.**
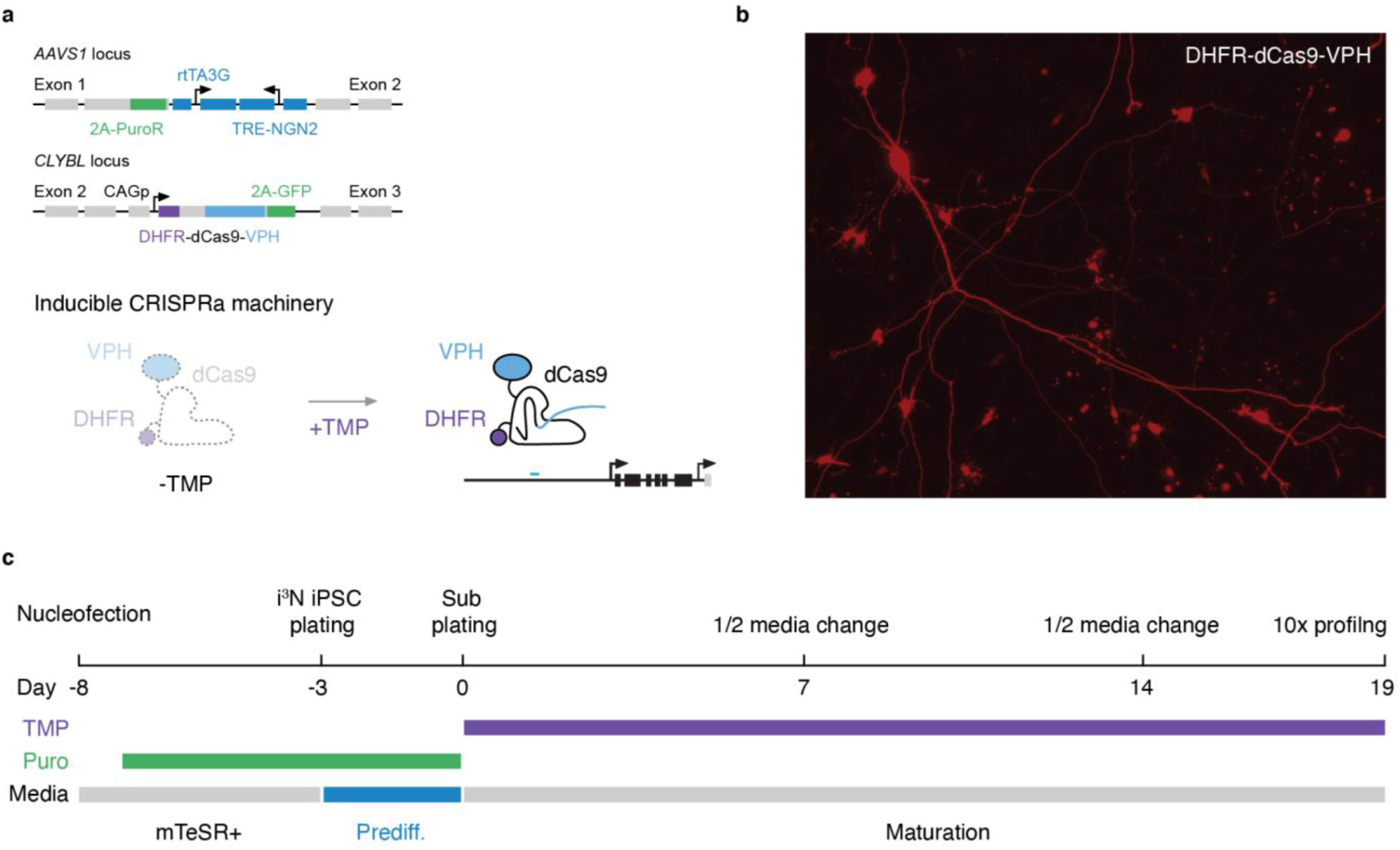
Inducible CRISPRa iPSC-derived neuron line functional validation, selection, and differentiation timeline. **a)** (Top) iPSCs equipped with a Dox-inducible *NGN2* transcription factor to drive neural differentiation (integrated at the *AAVS1* safe harbor locus) and TMP-inducible CRISPRa-VPH machinery (integrated at the *CLYBL* locus) were used for all iPSC-derived neuron experiments. (Bottom) In the absence of TMP, CRISPRa-VPH machinery is degraded via a DHFR degron. In the presence of TMP, the CRISPR-VPH machinery is stabilized, enabling perturbation. **b)** Functional validation of CRISPRa machinery in iPSC-derived neurons. Neurons were lipofected with a minP-tdTomato reporter and sgRNA that targets the minimal promoter. CRISPRa machinery drove clear tdTomato expression in differentiated neurons. **c)** Nucleofection, selection, and differentiation timeline. iPSCs were nucleofected with piggyFlex gRNA constructs at a high MOI and selected with puromycin to enrich cells for with multiple integrated gRNAs. Following differentiation induction neurons were subplated in maturation media with TMP to induce CRISPRa machinery. Neurons were single cell profiled following 19 days of differentiation (10x Genomics V3.1 chemistry with direct gRNA capture).

**Figure S7.**
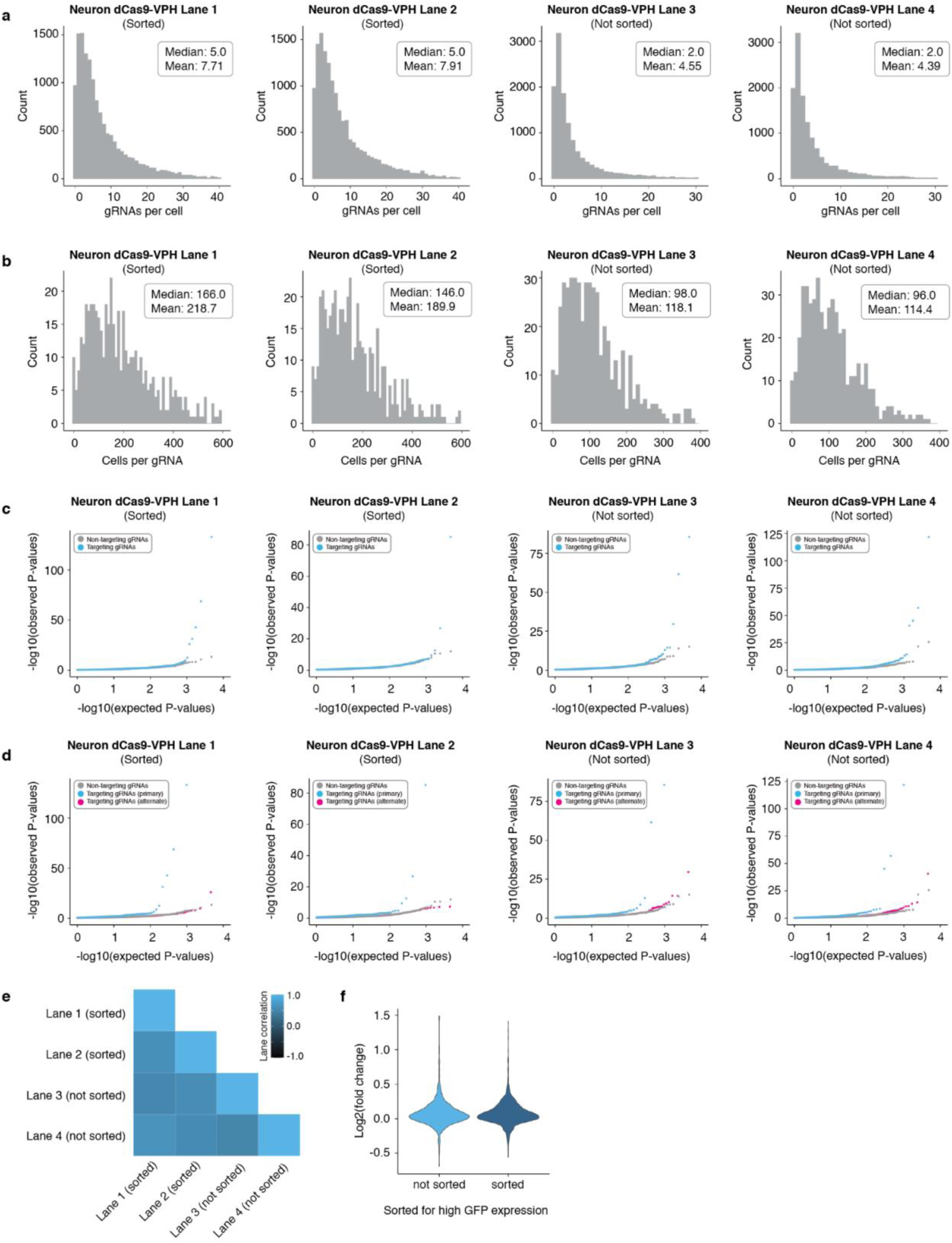
Results for four independent 10x Genomics lanes from iPSC-derived neuron screen. a) The four 10x Genomics lanes profiled consisted of two lanes with dCas9-VPH neurons that were sorted on the top 40% of GFP expression in these cells, and two lanes that were not on the top 40% of GFP expression in these cells. The cells that were not sorted were still 100% GFP+. Following QC and gRNA assignment we identified an average of 7.71, 7.91, 4.55, and 4.39 gRNAs/cell for the four different 10x Genomics lanes profiled (median 7.71, 7.91, 4.55, and 4.39 gRNAs per cell). Note that sorting neurons on the top 40% of GFP expression boosted the median and mean gRNAs/cell ∼2 fold. PiggyBac integrations per cell distribution is not well-modeled by a standard Poisson distribution and is better approximated by an exponential function. b) Multiplexing multiple perturbations per cell yielded an average of 218.7, 189.9, 118.1, and 114.4 cells/gRNA for the four different 10x Genomics lanes profiled (median 166, 146, 98, and 96 cells/gRNA). c) QQ-plots displaying observed vs. expected *P*-value distributions for targeting (blue) and NTC (downsampled) populations across the four different 10x Genomics lanes profiled. d) QQ-plots for targeting tests against their intended/programmed target (blue) compared to targeting tests of all other genes with 1Mb of each gRNA (pink) and NTCs (gray downsampled) across the four different 10x Genomics lanes profiled. e) Matrix correlation plot displaying the Pearson correlations of the log2 fold-changes of target gene expression values for programmed targets across the four different 10x Genomics lanes profiled. f) Violin plot displaying the log2 fold-changes of target gene expression values for programmed targets for neurons that were sorted on the top 40% GFP expression (sorted) and neurons that were not sorted (not sorted).

**Figure S8.**
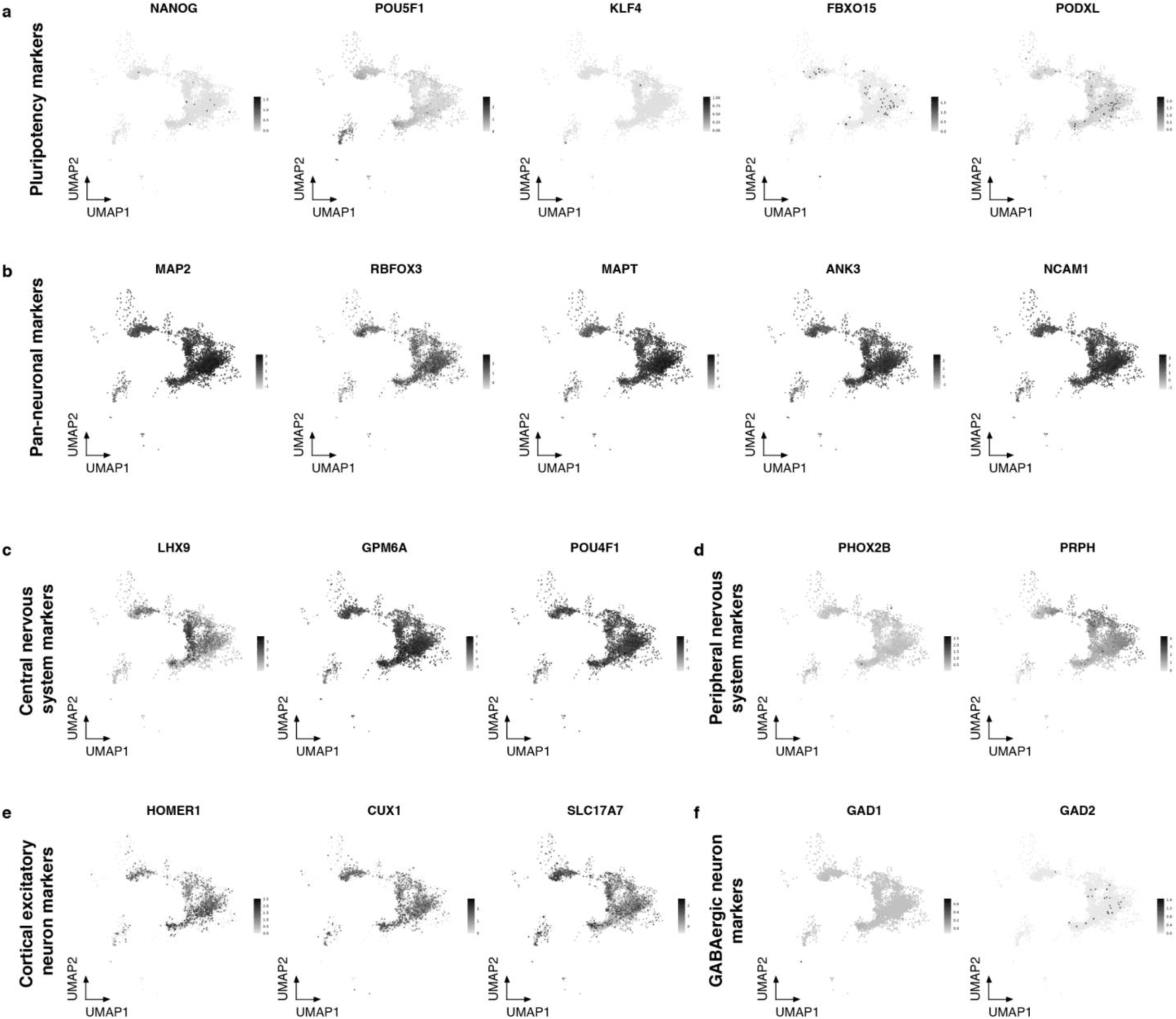
Single-cell transcriptomic characterization of iPSC-derived neurons used in screen. a) Expression feature plots of canonical pluripotency markers *NANOG*, *POU5F1*, *KLF4*, *FBXO15*, and *PODXL*. b) Expression feature plots of pan-neuronal markers *MAP2*, *RBFOX3*, *MAPT*, *ANK3*, and *NCAM1*. c) Expression feature plots of central nervous system marker genes *LHX9*, *GPM6A*, and *POU4F1*. d) Expression feature plots of peripheral nervous system marker genes *PHOX2B* and *PRPH*. e) Expression feature plots of cortical excitatory neuron markers *HOMER1*, *CUX1*, and *SLC17A7*. f) Expression feature plots of GABAergic neuron marker genes *GAD1* and *GAD2*.

**Figure S9.**
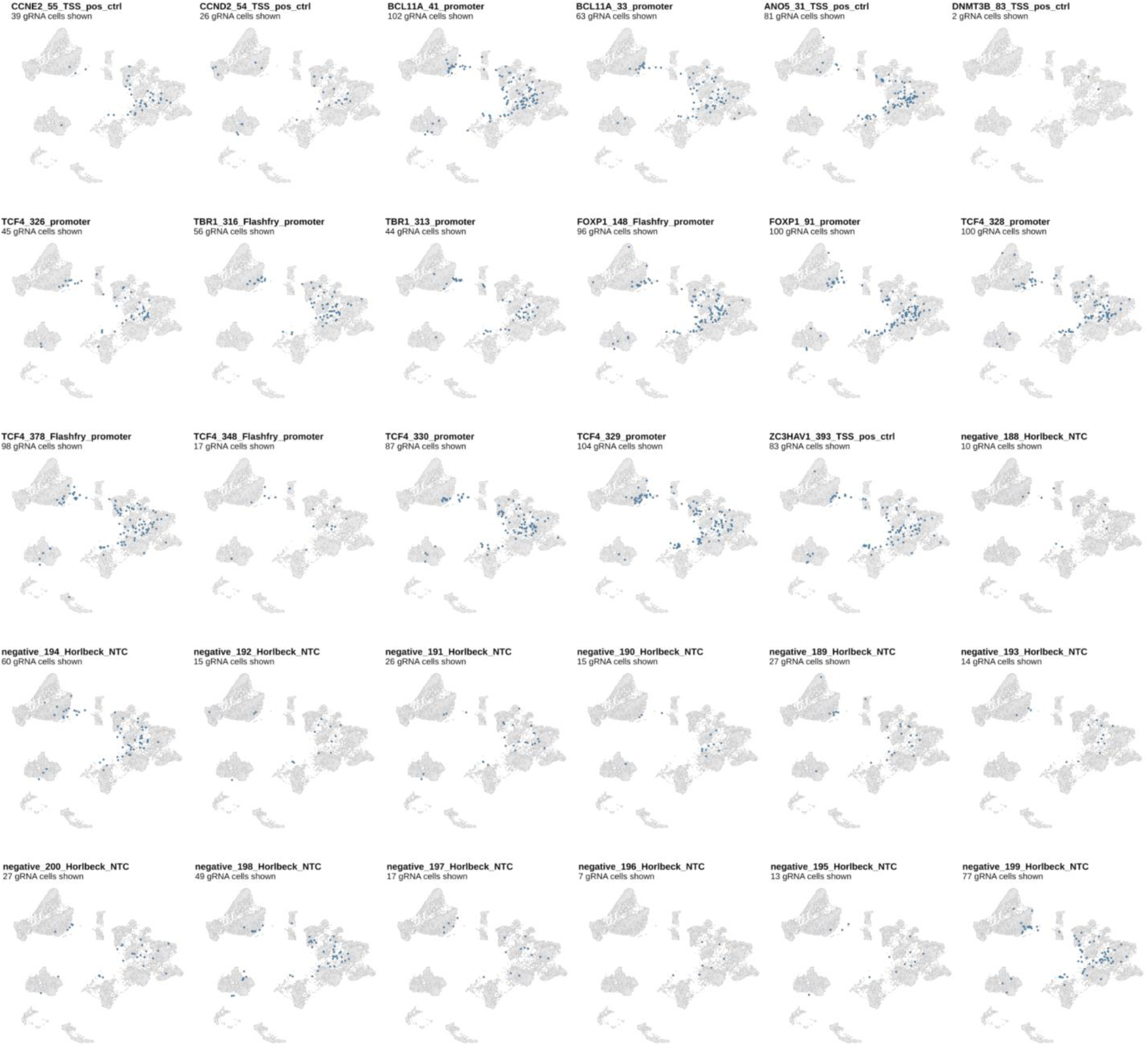
Distribution of CRISPRa gRNAs in single-cell neuron transcriptome data. Cells harboring specific CRISPRa gRNAs (dark blue) overlaid onto *NGN2*-induced neuron differentiation transcriptome data^28^. No readily apparent spatial enrichment of gRNAs is observed in UMAP plots. Note that the CRISPRa dataset was randomly downsampled to 5000 cells for all UMAP comparison analyses.

**Figure S10.**
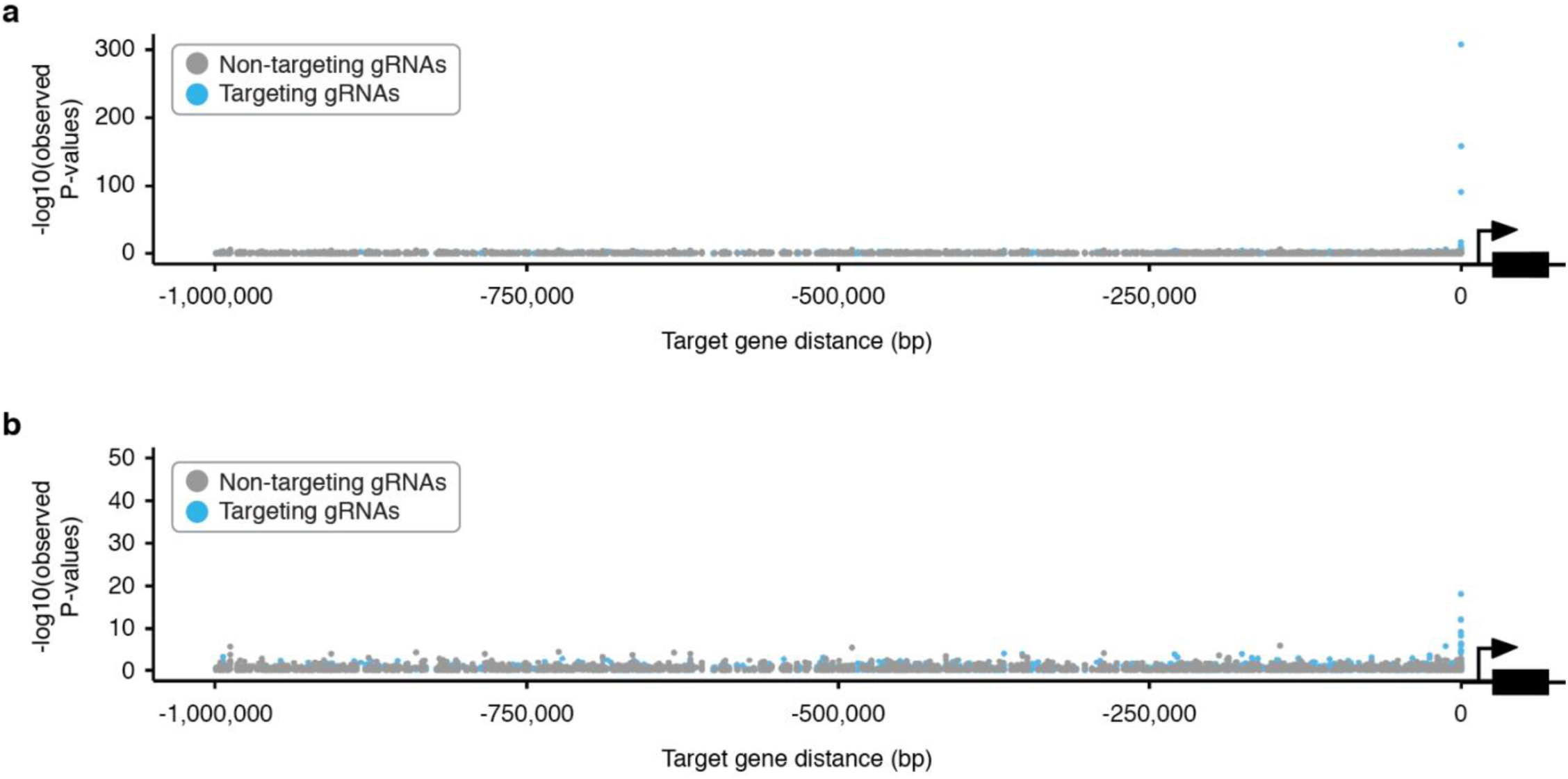
Successful targeting gRNAs are enriched for genomic proximity to their paired target gene scores near target genes in the iPSC-derived neurons. a) Targeting gRNAs yielding significant upregulation are enriched for proximity to their target gene, while NTCs are not. b) Same plot as in a, with the y-axis clipped at 50.

**Figure S11.**
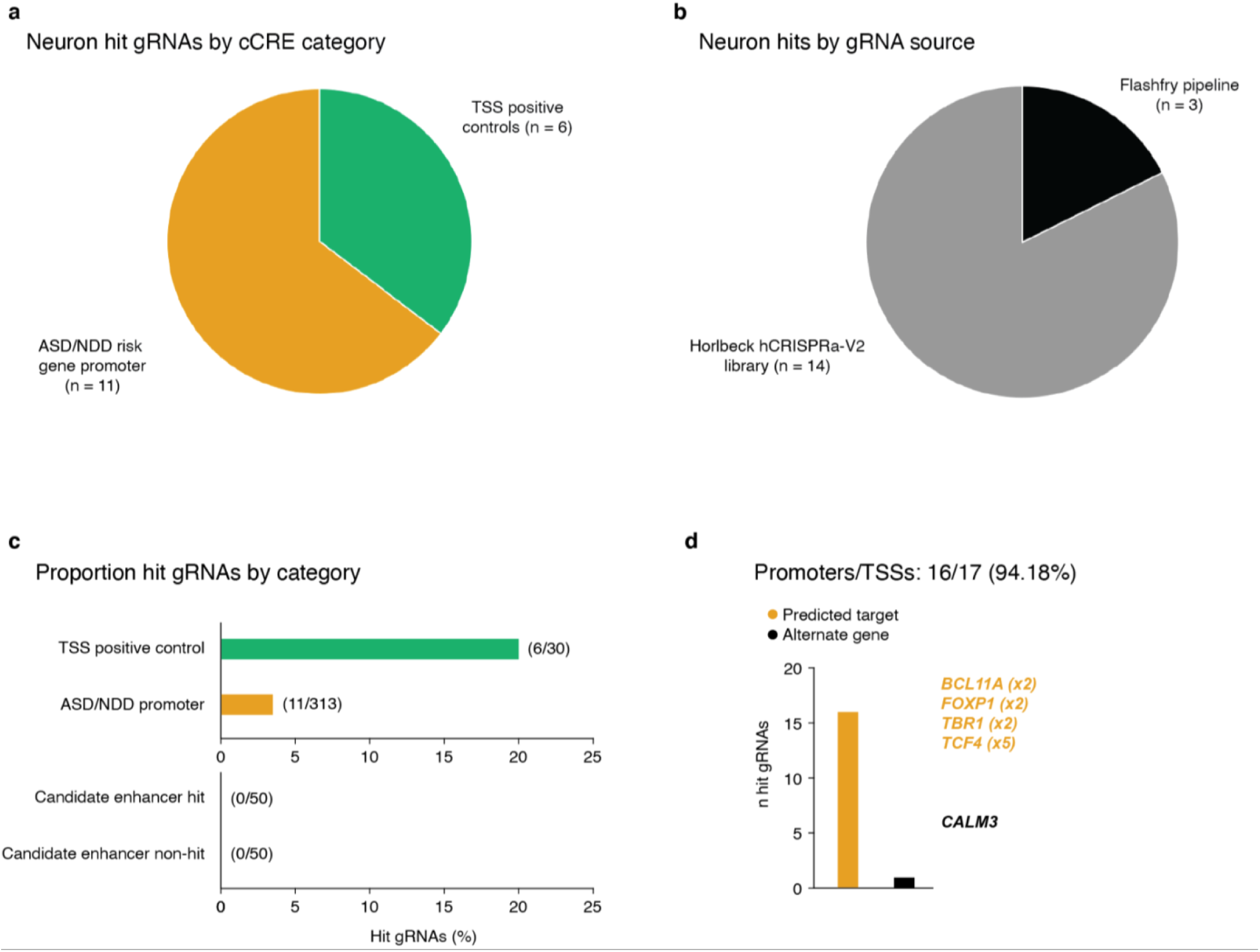
Hit breakdown for screen conducted in iPSC-derived neurons. a) Neuron hit gRNAs by cCRE category. b) Neuron hit gRNAs by gRNA source library or design pipeline. c) Proportion of hit gRNAs by cCRE category. d) Proportion of hit gRNAs yielding upregulation of their intended/expected target gene or an alternate gene for candidate promoters/TSSs. Example hits targeting candidate NDD risk gene promoters are listed. Bracketed numbers denote the number of independent hit gRNAs targeting the same cCRE.

**Figure S12.**
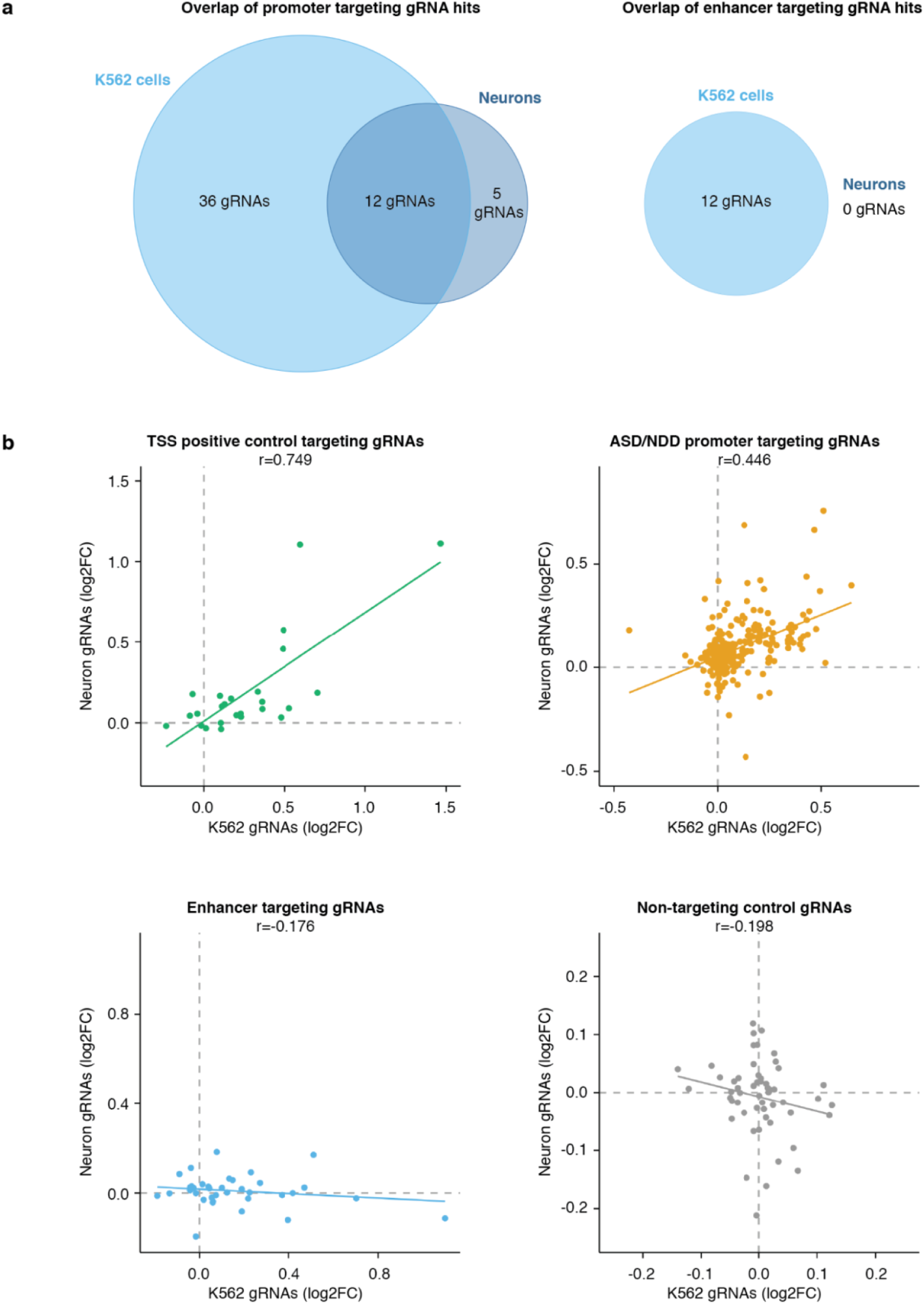
Comparison of K562 vs. neuronal CRISPRa screening hits. **a)** Venn diagram showing number of overlapping promoter-targeting gRNA hits (left) and enhancer-targeting gRNA hits (right) between the K562 and neuron CRISPRa screens. **b)** Correlation plots of log2 fold changes of TSS positive control targeting gRNAs (top left), ASD/NDD promoter targeting gRNAs (top right), enhancer targeting gRNAs (bottom left), and NTC gRNAs (bottom right) between the K562 and neuron CRISPRa screens.

**Figure S13.**
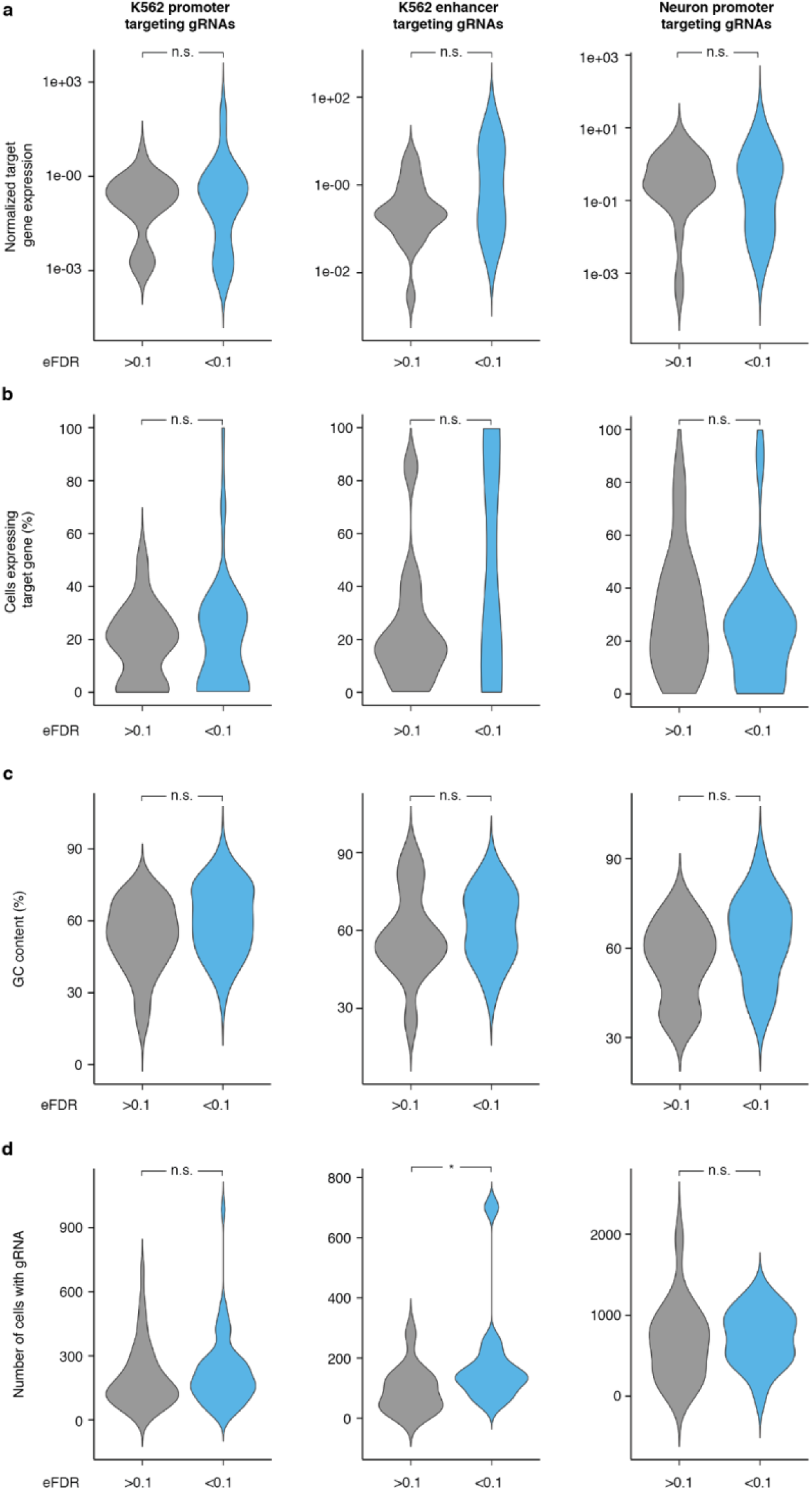
Characteristics of gRNAs leading to upregulation at EFDR<0.1vs. EFDR>0.1. a) Comparison of normalized gene expression values of targeted genes of gRNAs that resulted in an EFDR<0.1 (designated as “hit” gRNAs) versus gRNAs that resulted in an EFDR>0.1 (not designated as “hit” gRNAs). b) Comparison of the percentage of cells expressing the target gene of gRNAs that resulted in an EFDR<0.1 versus gRNAs that resulted in an EFDR>0.1. c) GC content (in percent) of gRNAs that resulted in an EFDR<0.1 versus gRNAs that resulted in an EFDR>0.1. d) Number of cells harboring each gRNA for gRNAs that resulted in an EFDR<0.1 versus gRNAs that resulted in an EFDR>0.1. For all panels, K562 promoter-targeting gRNAs (left), K562 enhancer-targeting gRNAs (middle), and neuron promoter-targeting gRNAs (right) are shown. Abbreviations: n.s.: not significant (p>0.05, Wilcoxon rank-sum test), *: p<0.05 (Wilcoxon rank-sum test).

**Figure S14.**
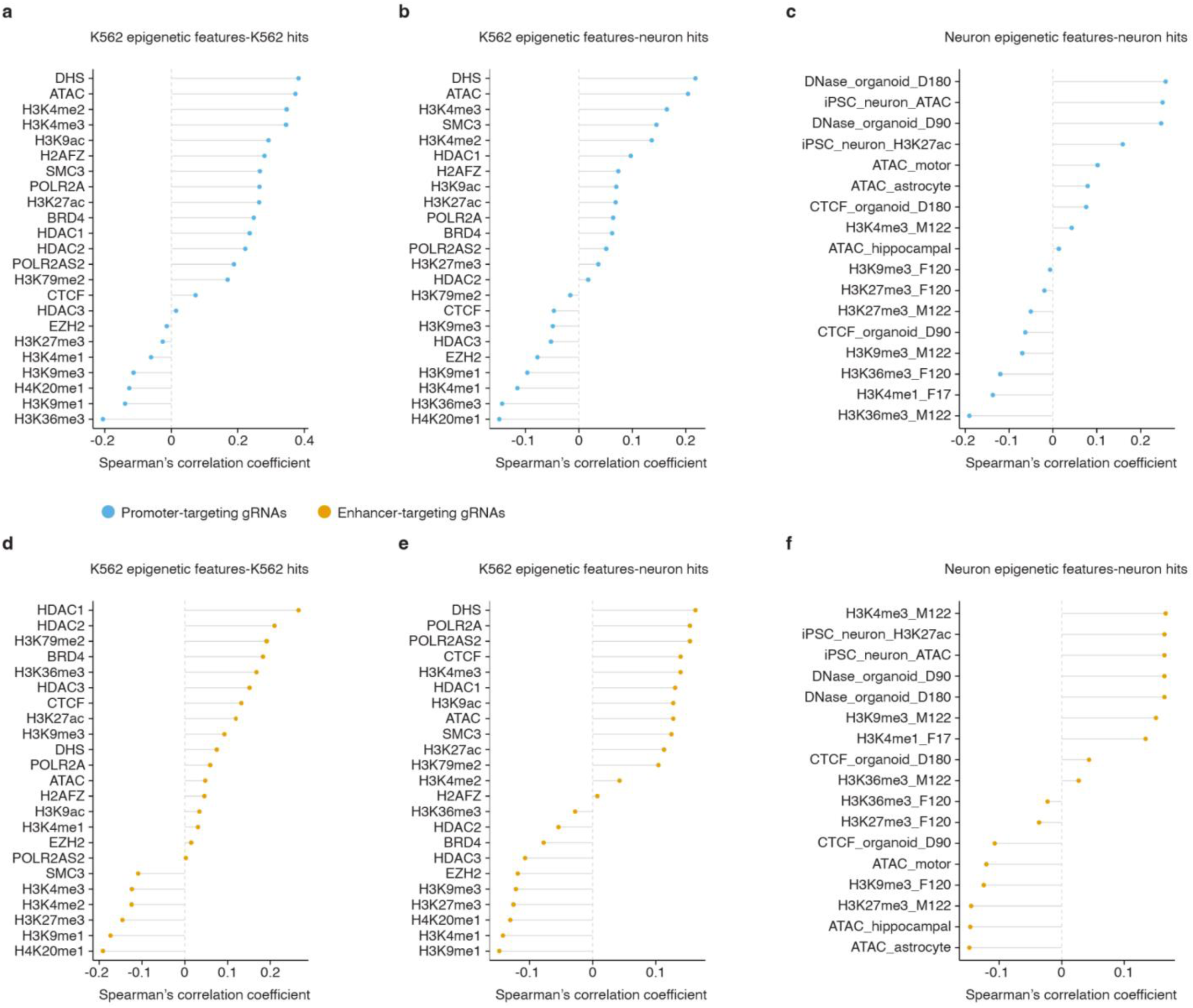
Epigenetic features predictive of CRISPRa hit gRNAs. a) Correlation of K562 epigenetic features with hit vs. non-hit status for promoter-targeting K562 gRNAs. b) Correlation of K562 epigenetic features with hit vs. non-hit status for promoter-targeting neuron gRNAs. c) Correlation of brain tissue and organoid epigenetic features with hit vs. non-hit status for promoter-targeting neuron gRNAs. d) Correlation of K562 epigenetic features with hit vs. non-hit status for enhancer-targeting K562 gRNAs. e) Correlation of K562 epigenetic features with hit vs. non-hit status for enhancer-targeting neuron hit gRNAs. f) Correlation of brain tissue and organoid epigenetic features with hit vs. non-hit status for enhancer-targeting neuron hit gRNAs. Although neuron hits are correlated with K562 epigenetic features, these correlations are lower in magnitude than for K562 hits; this pattern may follow from overlap in accessible vs. repressed regions between the cell types, together with the deep extent to which K562 cells have been epigenetically profiled.

**Figure S15.**
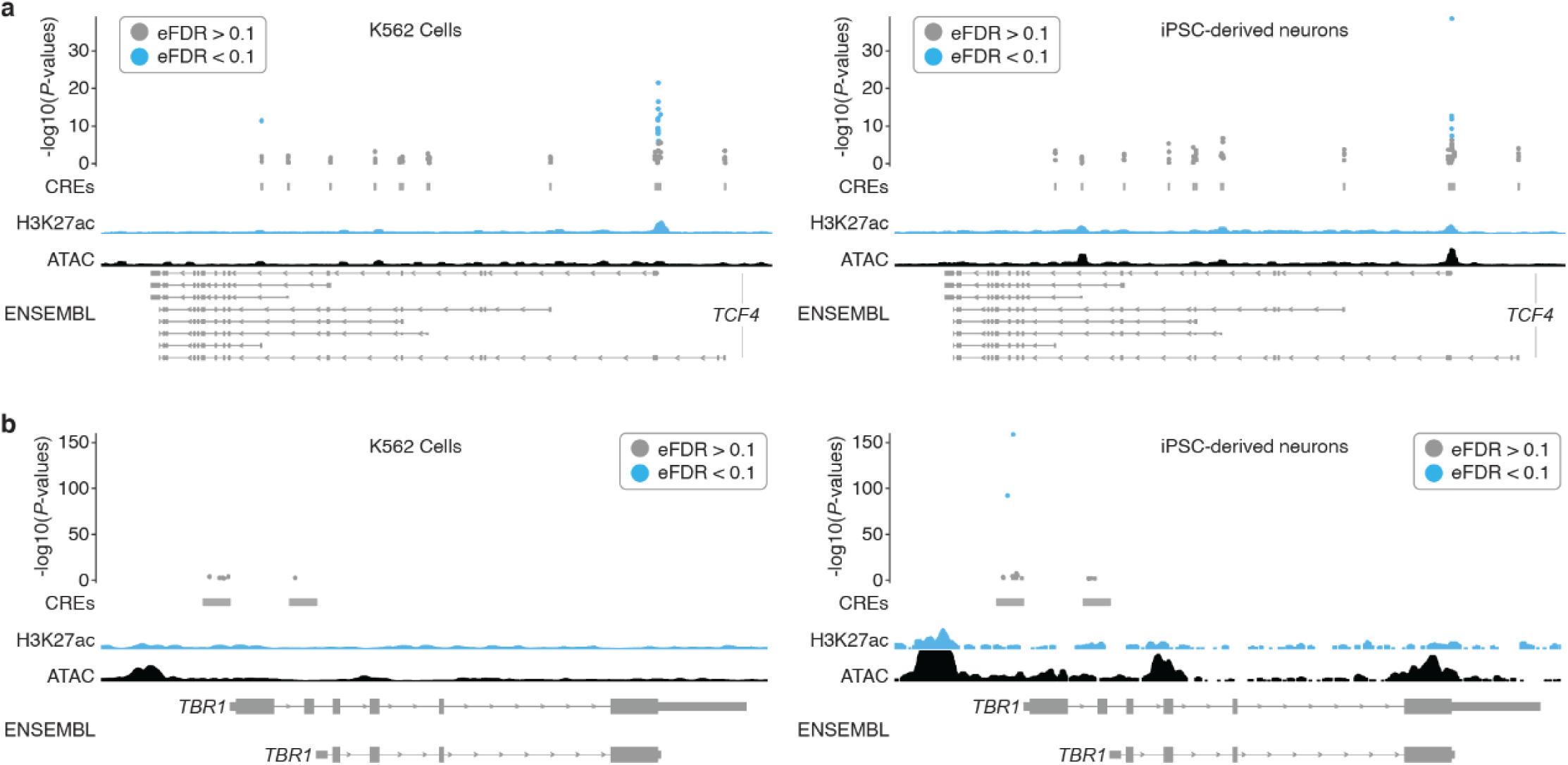
TSS and cell-type specific promoters. **a)** The majority of hit gRNAs for *TCF4* target the same TSS in K562 cells and iPSC-derived neurons. Empirical *P*-values are visualized alongside tracks for K562 ATAC-seq (ENCODE), K562 H3K27ac signal (ENCODE), iPSC-derived neuron ATAC-seq (accessibility)^45^, iPSC-derived neuron H3K27ac^45^ and RefSeq validated transcripts (ENSEMBL/NCBI). **b)** Two hit gRNAs targeting the same TSS of *TBR1* drive upregulation specifically in iPSC-derived neurons. Genomic tracks are the same as in panel **a**.

**Figure S16.**
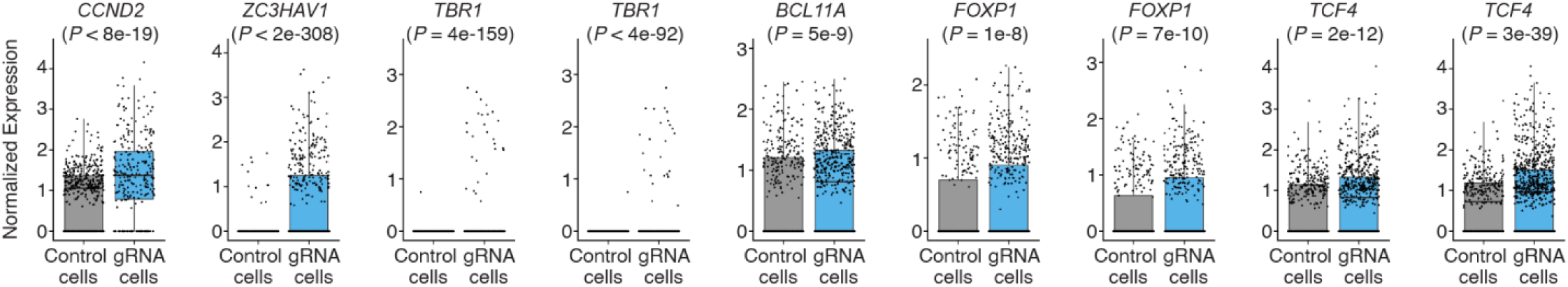
Example upregulations in iPSC-derived neurons. Example violin plots showing the average log2 fold change between cells with a given gRNA and controls for select hit gRNAs. Hits include TSS positive controls (*CCND2*, *ZC3HAV1*), candidate promoters of genes rarely or not expressed NGN2, including the cortical neuron marker *TBR1*, and candidate promoters of genes with native expression in iPSC-derived neurons that could be further upregulated (*BCL11A*, *FOXP1*, and *TCF4*). Number of cells bearing each targeting gRNA (from left to right): *CCND2* (n = 350), *ZC3HAV1* (n = 765), *TBR1* (n = 455), *TBR1* (n = 655), *BCL11A* (n = 964), *FOXP1* (n = 1198), *FOXP1* (n = 917), *TCF4* (n = 1051), *TCF4* (n = 1253). Control cells are downsampled to have the same number of cells as the average number of cells detected per gRNA (n = 638) for visualization. Normalized expression values represent log normalized expression values from Seurat.

**Figure S17.**
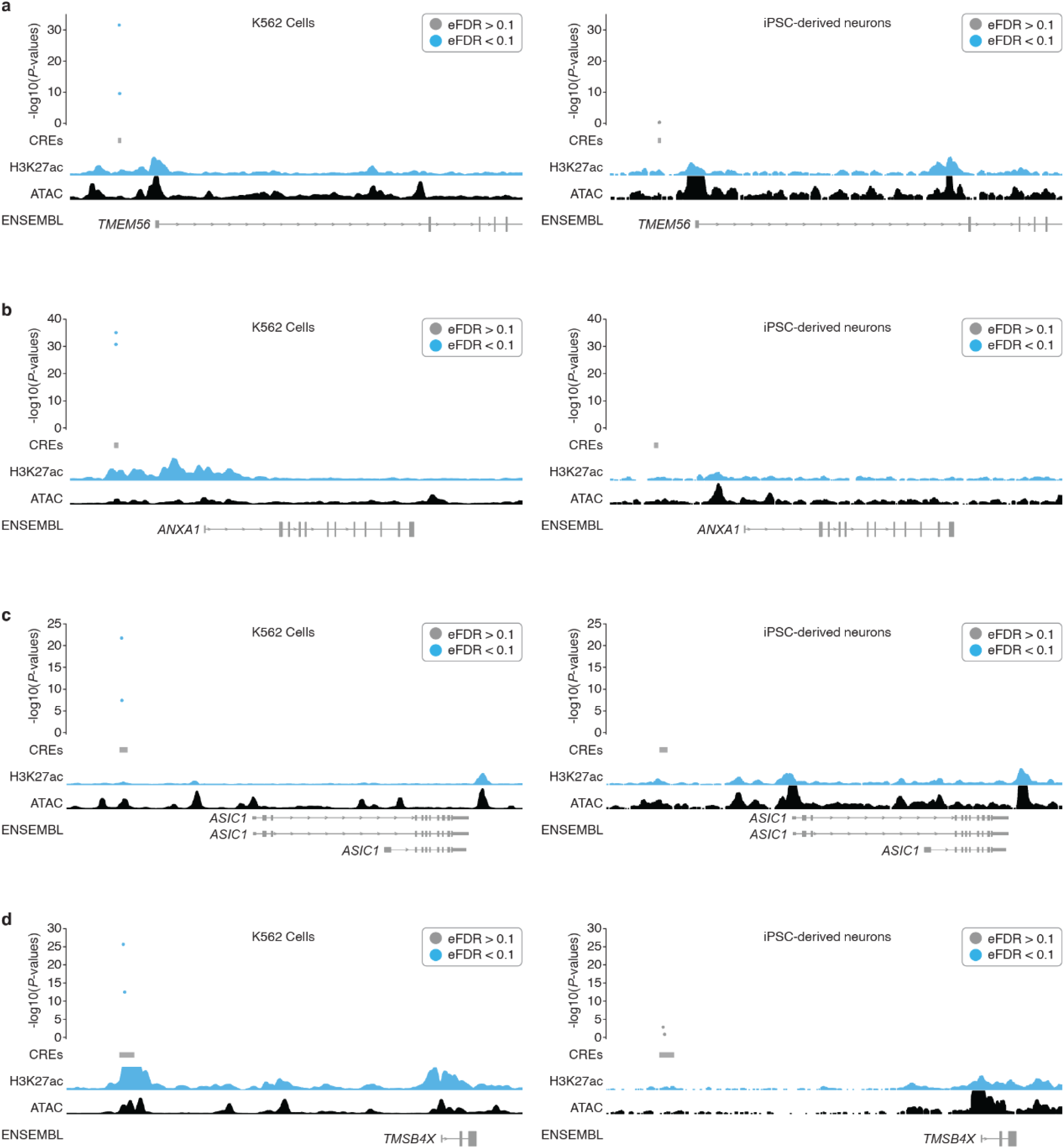
Cell-type specific enhancers. a-d) Empirical *P*-values are visualized alongside tracks for K562 ATAC-seq (ENCODE), K562 H3K27ac signal (ENCODE), iPSC-derived neuron ATAC-seq (accessibility)^45^, iPSC-derived neuron H3K27ac^45^ and RefSeq validated transcripts (ENSEMBL/NCBI). All K562 enhancer hits were cell type specific. Enhancers with multiple hit gRNAs are shown.

**Figure S18.**
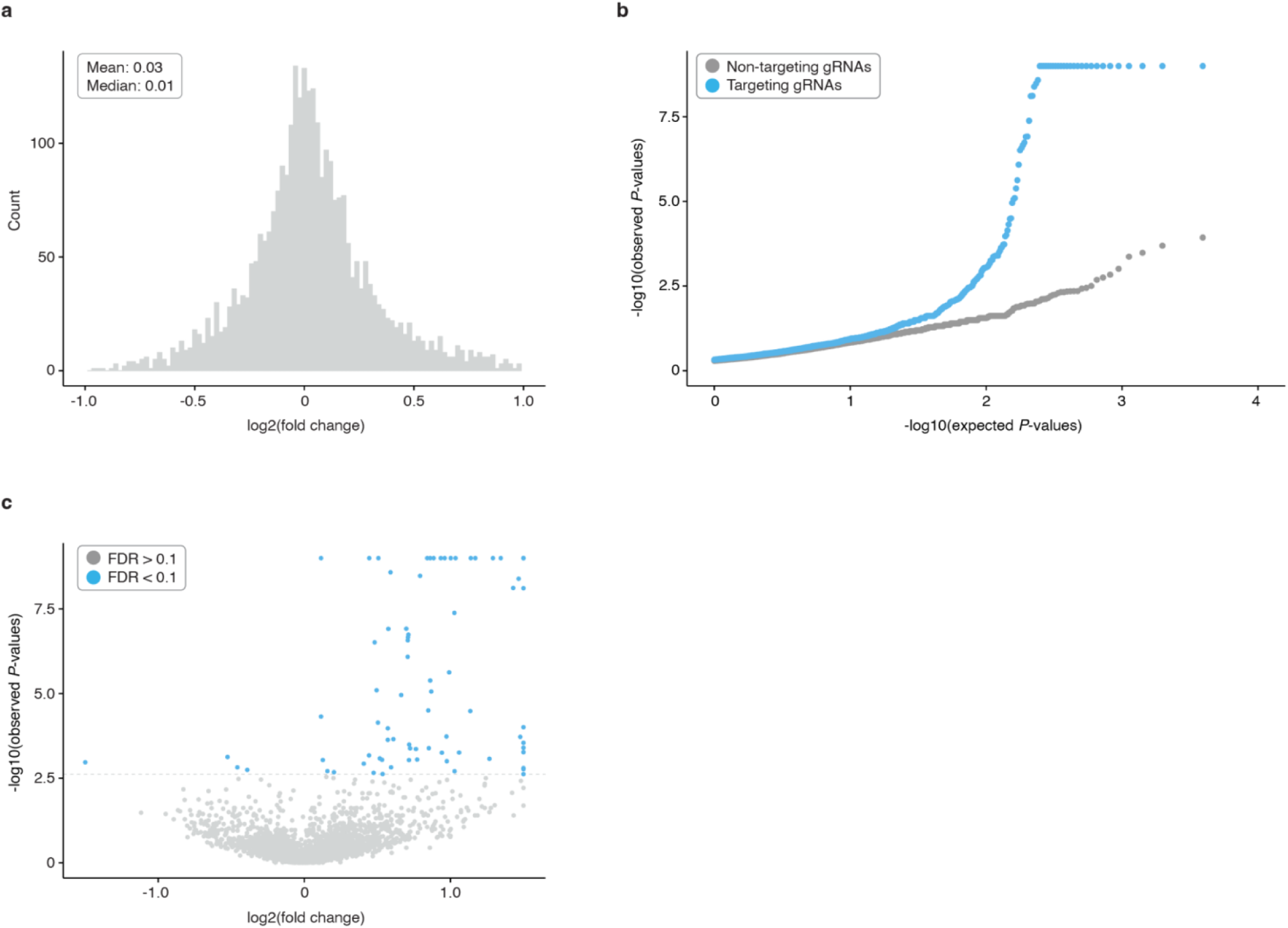
SCEPTRE results for K562 cells. a) Log2 fold change distribution for NTC gRNA tests in SCEPTRE framework. A NTC distribution centered at zero is expected for well-calibrated screens. b) QQ-plot showing the distribution of expected vs. observed *P*-values for targeting (blue) and non-targeting (gray, downsampled) differential expression tests from SCEPTRE. There is an excess of highly significant *P*-values in targeting tests compared to controls. c) Volcano plot showing the average log2 fold change and *P*-values for targeting tests.

**Figure S19.**
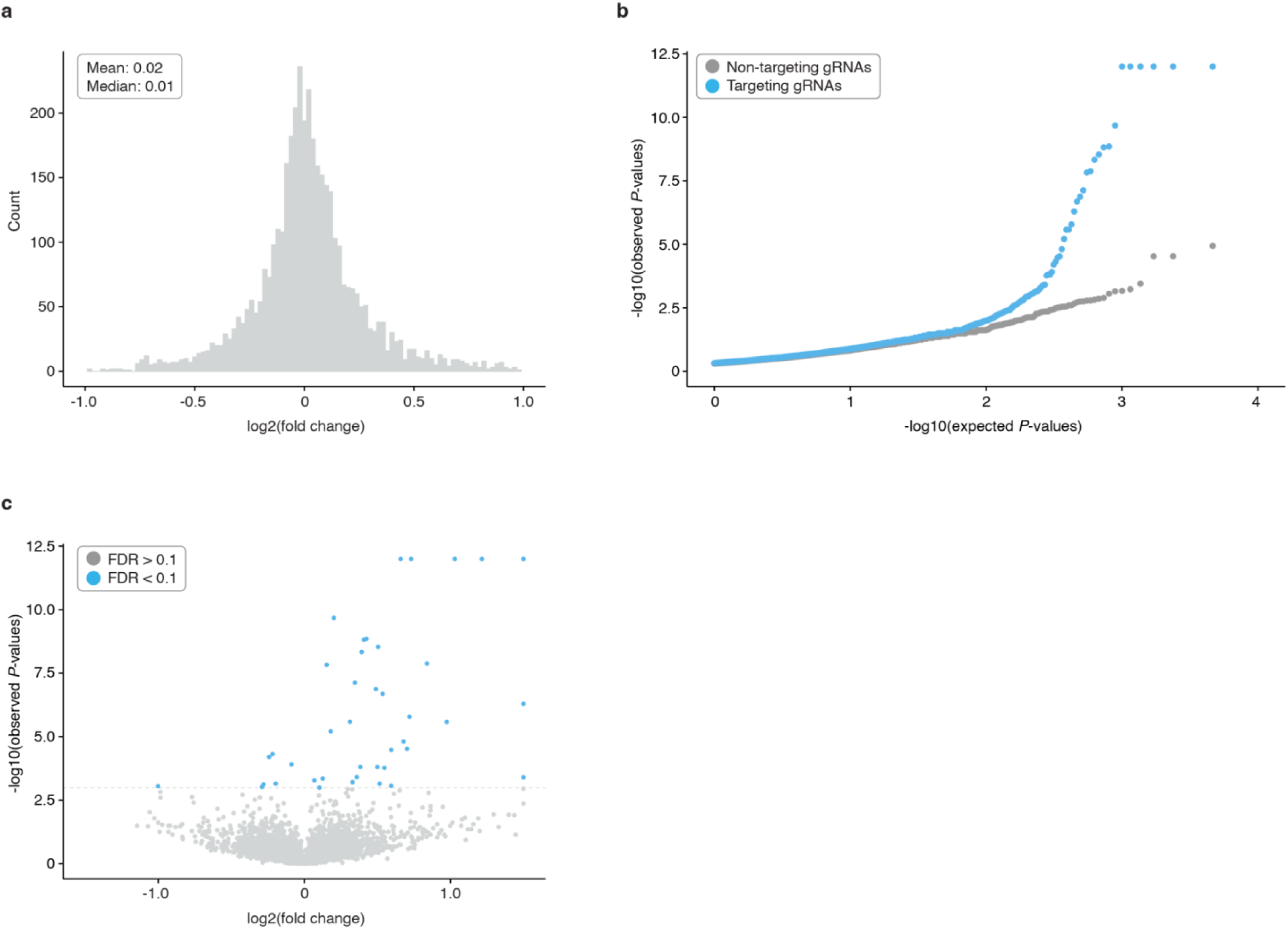
SCEPTRE results for iPSC-derived neurons. a) Log2 fold change distribution for NTC gRNA tests in SCEPTRE framework. A NTC distribution centered at zero is expected for well-calibrated screens. b) QQ-plot showing the distribution of expected vs. observed *P*-values for targeting (blue) and non-targeting (gray, downsampled) differential expression tests from SCEPTRE. There is an excess of highly significant *P*-values in targeting tests compared to controls. c) Volcano plot showing the average log2 fold change and *P*-values for targeting tests.

**Figure S20.**
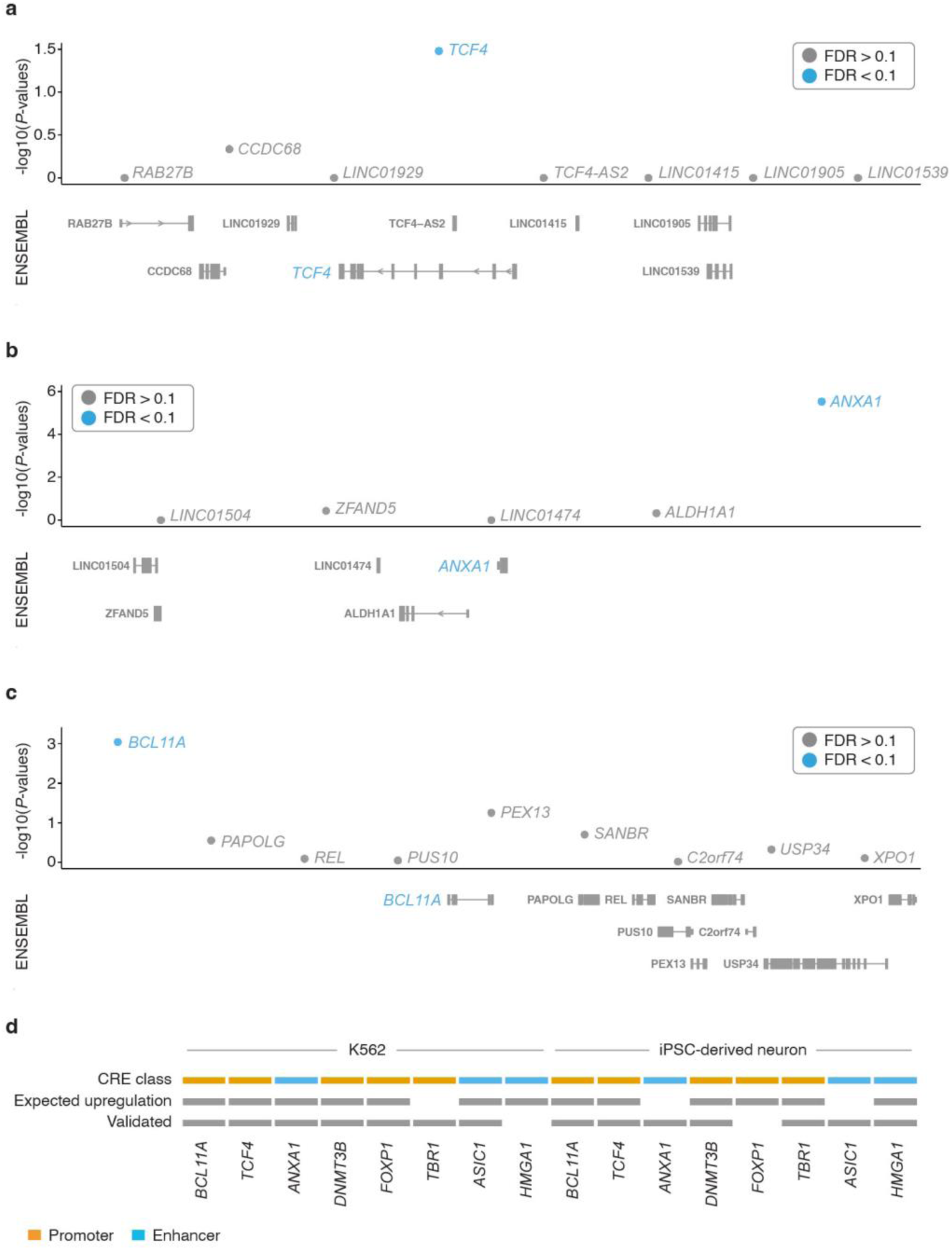
Singleton validations confirm target upregulations. **a)** *TCF4* upregulation in K562 cells expressing a *TCF4* promoter-targeting gRNA compared to control cells. *P*-values (DESeq2) from differential expression tests of the intended target (blue) are visualized alongside neighboring genes within 1Mb of the target site (gray). Neighboring genes are filtered to those detected in the sc-RNA-seq data to aid visualization. **b)** *ANXA1* upregulation in K562 cells expressing an *e-ANXA1* enhancer-targeting gRNA compared to control cells. Genomic tracks are the same as in panel **a**. **c)** *BCL11A* upregulation in iPSC-derived neurons expressing an *BCL11A* promoter-targeting gRNA compared to control cells. Genomic tracks are the same as in panel **a**. **d)** Categorical heatmap indicating whether a given validation gRNA targeted a promoter or enhancer, whether the gRNA was expected to drive upregulation in that cell type based on the multiplex screen, and whether the expected upregulation (or not) was observed in singleton validations.

**Figure S21.**
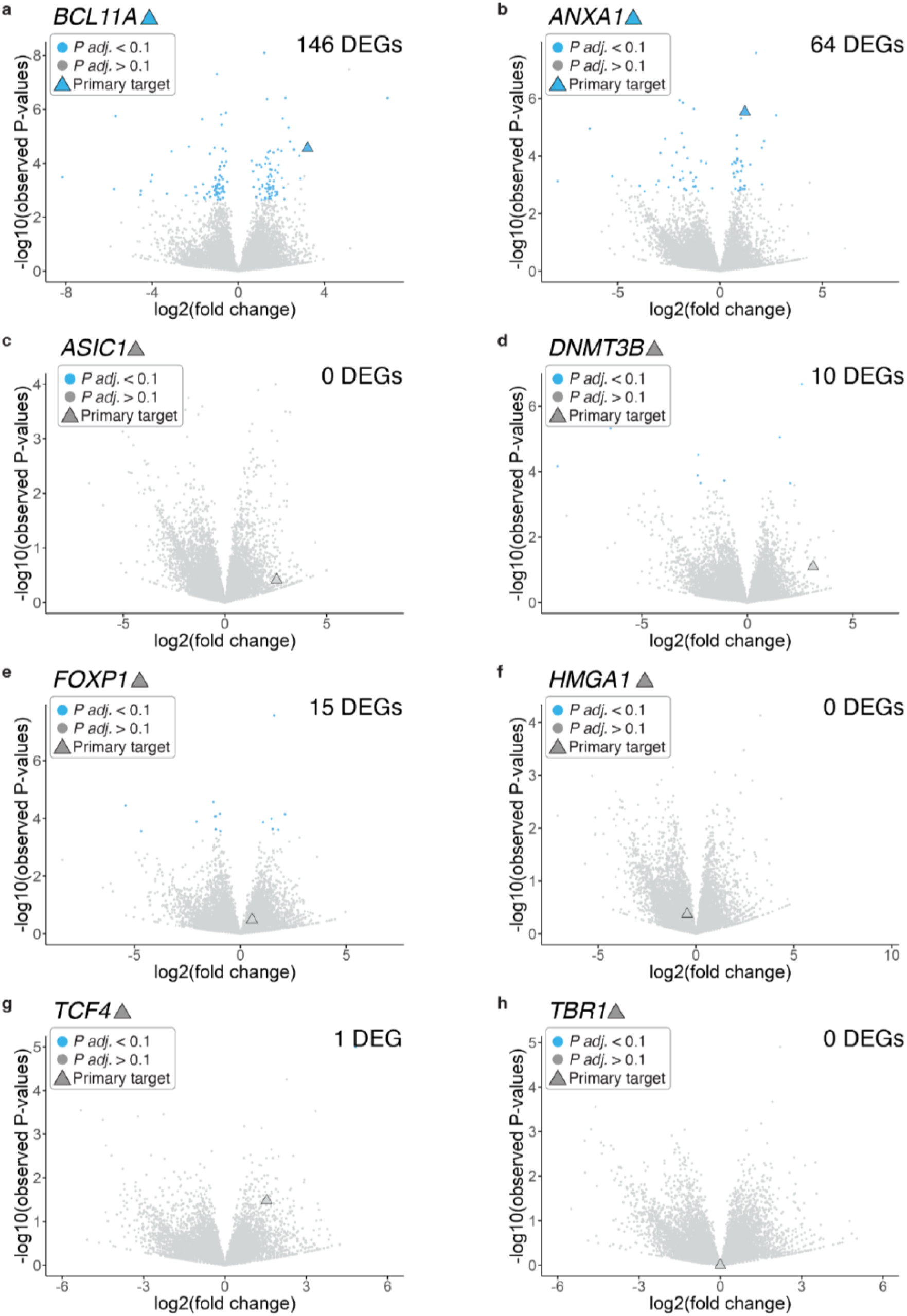
Genome-wide differential expression analysis of singleton validations in K562cells. a) Genome-wide differential gene expression analysis comparing cell lines harboring a *BCL11A* promoter-targeting gRNA compared to all other lines as a control. b-h) Differential gene expression analyses as in panel a) for gRNAs targeting cCREs of: *ANXA1* (b), *ASIC1* (c), *DNMT3B* (d), *FOXP1* (e), HMGA1 (f), TCF4 (g), and TBR1 (h). Differential expression *P*-values and number of Differentially Expressed Genes (DEGs) from DESeq2.

**Figure S22.**
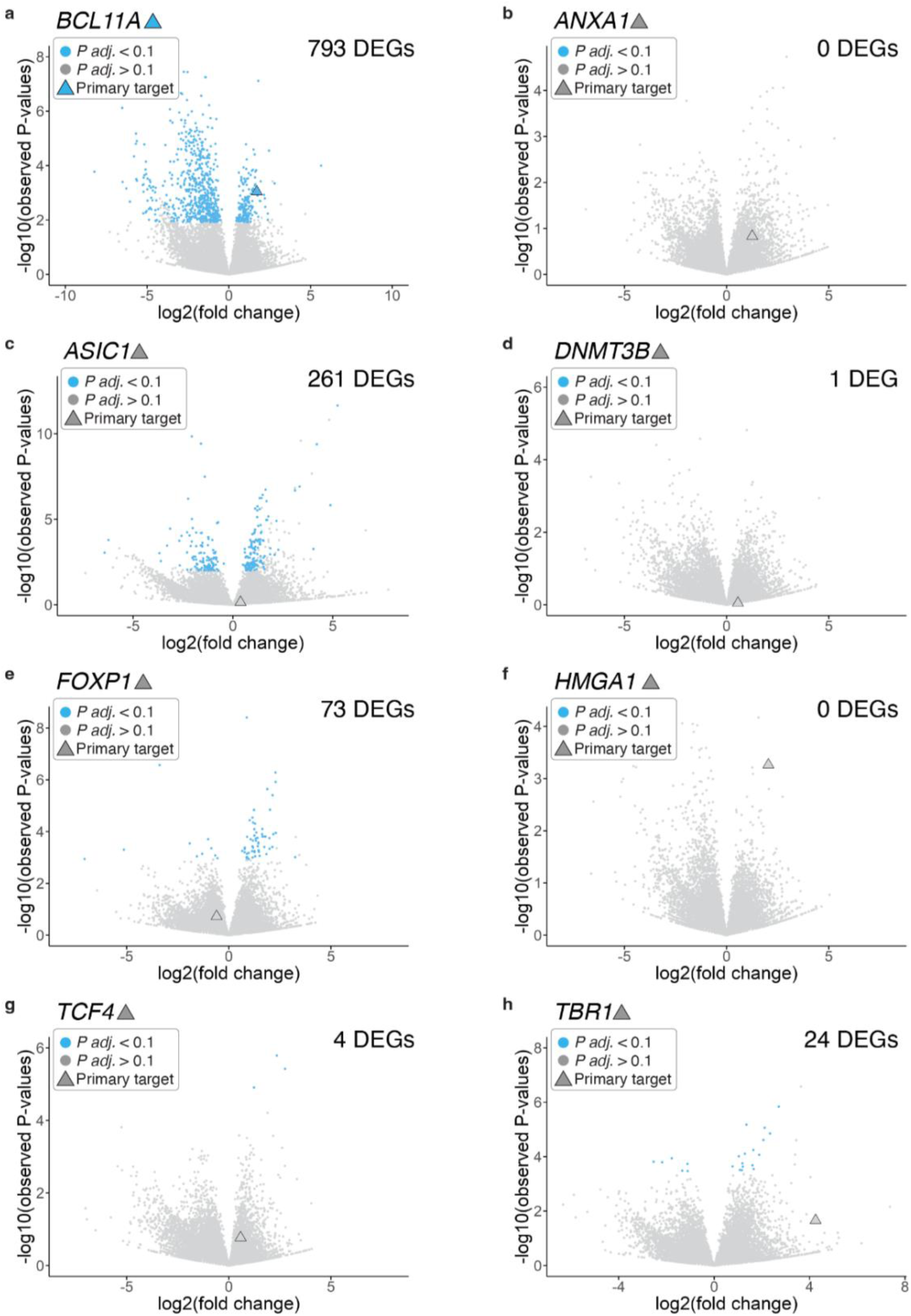
Genome-wide differential expression analysis of singleton validations in iPSC-derived neurons. a) Genome-wide differential gene expression analysis comparing cell lines harboring a *BCL11A* promoter-targeting gRNA compared to all other lines as a control. b-h) Differential gene expression analyses as in panel a) for gRNAs targeting cCREs of: *ANXA1* (b), *ASIC1* (c), *DNMT3B* (d), *FOXP1* (e), HMGA1 (f), TCF4 (g), and TBR1 (h). Differential expression *P*-values and number of Differentially Expressed Genes (DEGs) from DESeq2.

**Figure S23.**
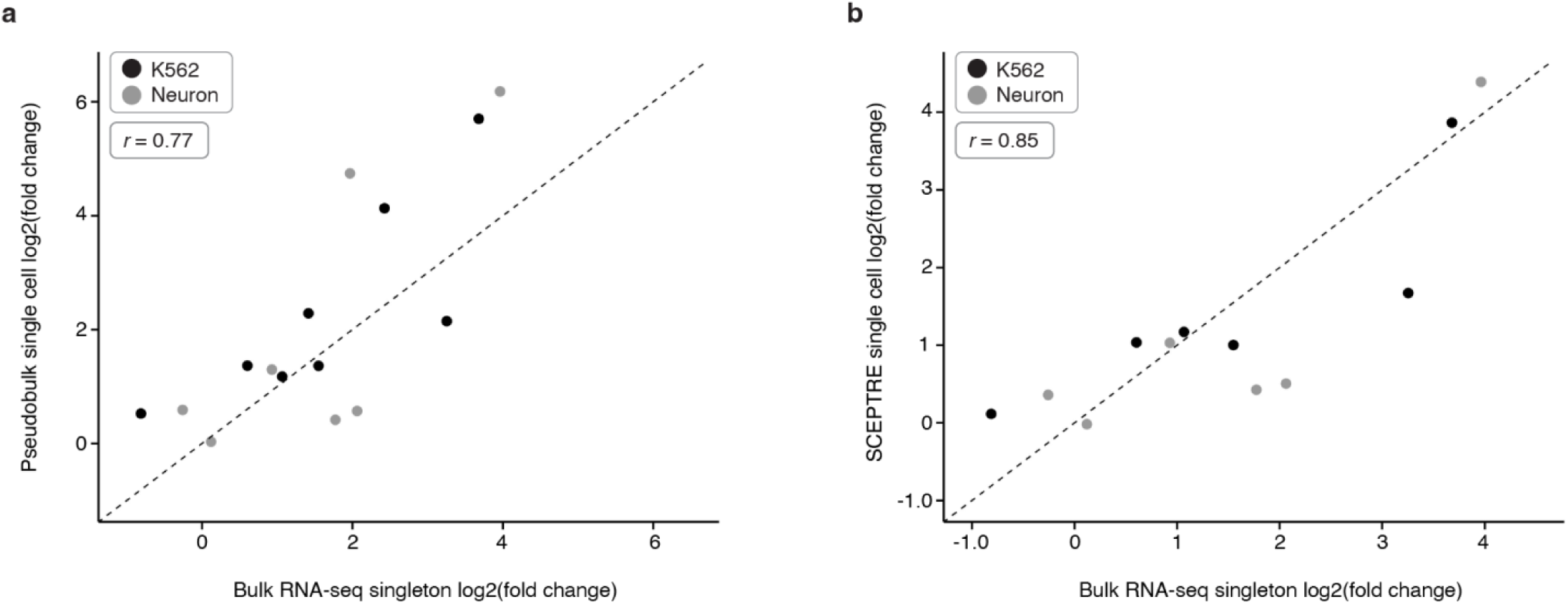
Correlation of fold change estimates from singleton bulk RNA-seq and multiplex single cell CRISPRa experiments. **a)** Correlation of log2 fold change estimates from singleton bulk RNA-seq CRISPRa experiments and pseudobulked single cells from the multiplex single cell CRISPRa experiment. Note that TBR1 expression was not detected in K562 cells precluding calculation of fold changes. Points are coloured according to cell context (K562 or iPSC-derived neurons). **b)** Correlation of log2 fold change estimates from singleton bulk RNA-seq CRISPRa experiments and SCEPTRE log2 fold change estimates from the multiplex single cell CRISPRa experiment. Note that *TBR1* and *ASIC1* in K562, and *DNMT3B* and *ANXA1* in iPSC-derived neurons, did not meet SCEPTRE’s minimum threshold for proportion of cells expressing a target gene, precluding calculation of fold changes. Points are coloured according to cell context (K562 or iPSC-derived neurons). Modestly increased correlation between SCEPTRE and bulk log2 fold change estimates is potentially due to accounting for various technical factors (sequencing depth, cell quality, etc.) that direct pseudobulking does not. Pearson’s correlation coefficient is shown.

## Acknowledgements

We are grateful to members of the Shendure and Ahituv labs, as well as members of the Sanders, Bender, Bateup, and Feldman labs for comments, suggestions, and discussions on this work. We are particularly grateful to the Shendure lab gene regulation subgroup for technical advice and deep discussions regarding the development of the CRISPRa screening method. We would also like to thank Haedong Kim who piloted the in-house bulk RNA-seq protocol and provided reagents for the bulk RNA-seq experiments. The Human WTC11 *NGN2* ecDHFR-dCas9-VPH line was a kind gift from the M. Kampmann lab at UCSF. Lenti dCAS9-VP64_GFP (Addgene plasmid # 61422) was a kind gift from the F. Zhang lab at Broad/MIT.

## Author contributions

Conceptualization, J.S. and N.A.; Investigation, F.M.C., T.A.M., N.F.P., and R.M.D.; Data Curation, F.M.C., T.A.M., N.F.P, and R.M.D.; Formal Analysis, F.M.C., T.A.M., and N.F.P.; Visualization, F.M.C. and T.A.M.; Resources, N.A. and J.S.; Supervision, L.S., S.J.S., N.A. and J.S.; Writing – Original Draft, F.M.C., T.A.M., J.S.; Writing – Review & Editing, F.M.C., T.A.M., N.F.P., B.M., S.D., S.R., J.B.L., D.C., R.M.D., L.S., S.J.S., N.A, and J.S.; Funding Acquisition, S.J.S., N.A., and J.S.

## Data Availability

Raw sequencing data and processed files generated in this study will be deposited to GEO. Raw data, processed data, code, and scripts used for analyses are all publicly available and are accessible via the following website: https://krishna.gs.washington.edu/content/members/CRISPRa_QTL_website/public/.

## Supplementary Materials

### Figures S1-S21

1. **Figure S1** - gRNA design pipeline, library contents, and piggyFlex gRNA delivery construct.
2. **Figure S2** - Functional validation of CRISPRa K562 cell lines.
3. **Figure S3** - Results for four independent 10x Genomics lanes from K562 screen.
4. **Figure S4** - Hit breakdown for screen conducted in K562 cells.
5. **Figure S5** - Example upregulations in K562 cells.
6. **Figure S6** - Inducible CRISPRa iPSC-derived neuron line functional validation, selection, and differentiation timeline.
7. **Figure S7** - Results for four independent 10x Genomics lanes from iPSC-derived neuron screen.
8. **Figure S8** - Single-cell transcriptomic characterization of iPSC-derived neurons used in screen.
9. **Figure S9** - Distribution of CRISPRa gRNAs in single-cell neuron transcriptome data.
10. **Figure S10** - Successful targeting gRNAs are enriched for genomic proximity to their paired target gene scores near target genes in the iPSC-derived neurons.
11. **Figure S11** - Hit breakdown for screen conducted in iPSC-derived neurons.
12. **Figure S12** - Comparison of K562 vs. neuronal CRISPRa screening hits.
13. **Figure S13** - Characteristics of gRNAs leading to upregulation at EFDR<0.1 vs. EFDR>0.1.
14. **Figure S14** - Epigenetic features predictive of CRISPRa hit gRNAs.
15. **Figure S15** - TSS and cell-type specific promoters.
16. **Figure S16** - Example upregulations in iPSC-derived neurons.
17. **Figure S17** - Cell-type specific enhancers.
18. **Figure S18** - SCEPTRE results for K562 cells.
19. **Figure S19** - SCEPTRE results for iPSC-derived neurons.
20. **Figure S20** - Singleton validations confirm target upregulations.
21. **Figure S21** - Genome-wide differential expression analyses of singleton validations in K562 cells.
22. **Figure S22** - Genome-wide differential expression analyses of singleton validations in iPSC-derived neurons.
23. **Figure S23** - Correlation of fold change estimates from singleton bulk RNA-seq and multiplex single cell CRISPRa experiments.

### Tables S1-S15

1. **Table S1** - gRNA sequences.
2. **Table S2** - K562 full screen results.
3. **Table S3** - K562 primary target results.
4. **Table S4** - K562 hits (EFDR < 0.1).
5. **Table S5** - iPSC-derived neuron full screen results.
6. **Table S6** - iPSC-derived neuron primary target results
7. **Table S7** - iPSC-derived neuron hits (EFDR < 0.1).
8. **Table S8** - Epigenetic feature datasets.
9. **Table S9** - SCEPTRE K562 NTC calibration results.
10. **Table S10** - SCEPTRE K562 targeting discovery results.
11. **Table S11** - SCEPTRE K562 hits (FDR < 0.1).
12. **Table S12** - SCEPTRE iPSC-derived neuron NTC calibration results.
13. **Table S13** - SCEPTRE iPSC-derived neuron targeting discovery results.
14. **Table S14** - SCEPTRE iPSC-derived neuron hits (FDR < 0.1).
15. **Table S15** - gRNA set for singleton validations.
16. **Table S16** - Differentially expressed genes from singleton validation experiments.

## Methods

### Cell Lines and Culture

#### K562 cell culture

K562s cells are a pseudotriploid ENCODE Tier I erythroleukemia cell line derived from a female (age 53) with chronic myelogenous leukemia^15^. All K562 cells were grown at 37°C, and cultured in RPMI 1640 + L-Glutamine (GIBCO, Cat. No. 11-875-093) supplemented with 10% fetal bovine serum (Fisher Scientific, Cat No. SH3039603) and 1% penicillin-streptomycin (Thermo Fisher Scientific, Cat. No. 15070063).

#### Induced pluripotent stem cell (iPSC) culture

Human WTC11 iPSCs equipped with a doxycycline-inducible *NGN2* transgene expressed from the *AAVS1* safe-harbor locus as well as an ecDHFR-dCas9-VPH construct (VPH consists of 12 copies of VP16, fused with a P65-HSF1 activator domain) expressed from the CLYBL safe-harbor locus were a gift from the Kampmann lab^6^. These iPSCs were cultured in mTeSR Plus Basal Medium (Stemcell technologies; Cat. No. 100-0276) on Greiner Cellstar plates (Sigma-Aldrich; assorted Cat. Nos.) coated with Geltrex™ LDEV-Free, hESC-Qualified, Reduced Growth Factor Basement Membrane Matrix (Gibco; Cat. No. A1413302) diluted 1:100 in Knockout DMEM (GIBCO/Thermo Fisher Scientific; Cat. No. 10829018). mTeSR Plus Basal Medium was replaced every other day. When 70–80% confluent, cells were passaged by aspirating media, washing with DPBS (GIBCO/Thermo Fisher Scientific; Cat. No. 14190144), incubating with StemPro Accutase Cell Dissociation Reagent (GIBCO/Thermo Fisher Scientific; Cat. No. A1110501) at 37 °C for 5 min, diluting Accutase 1:1 in mTeSR Plus Basal Medium, collecting cells in conical tubes, centrifuging at 800*g* for 3 min, aspirating supernatant, resuspending cell pellet in mTeSR Plus Basal Medium supplemented with 0.1% dihydrochloride ROCK Inhibitor (Stemcell technologies; Cat. No. Y-27632), counting and plating onto Geltrex-coated plates at the desired number.

#### Human iPSC-derived neuronal cell culture, differentiation, and CRISPRa induction

The iPSCs described above were used for the differentiation protocol below. On day −3, iPSCs were dissociated and centrifuged as above, and pelleted cells were resuspended in Pre-Differentiation Medium containing the following: Knockout DMEM/F-12 (GIBCO/Thermo Fisher Scientific; Cat. No. 12660012) as the base, 1X MEM Non-Essential Amino Acids (GIBCO/Thermo Fisher Scientific; Cat. No. 11140050), 1X N-2 Supplement (GIBCO/ Thermo Fisher Scientific; Cat. No. 17502048), 10 ng/mL NT-3 (PeproTech; Cat. No. 450-03), 10ng/mL BDNF (PeproTech; Cat. No. 450-02), 1 ug/mL Laminin mouse protein (Thermo Fisher Scientific; Cat. No. 23017015), 10 nM ROCK inhibitor, and 2 mg/mL doxycycline hyclate (Sigma-Aldrich; Cat. No. D9891) to induce expression of *NGN2*. iPSCs were counted and plated at 800K cells per Geltrex-coated well of a 12-well plate in 1 mL of Pre-Differentiation Medium, for three days. At day −2 and day −1, media changes were performed using pre-differentiation medium without ROCK inhibitor. On day −1, 12-well plates for differentiation were coated with 15 ug/mL Poly-L-Ornithine (Sigma-Aldrich; Cat. No. P3655) in DPBS, and incubated overnight at 37 degrees Celsius. On day 0, the Poly-L-Ornithine coated plates were washed three times using DPBS, and the plates were air dried in a 37 degree Celsius incubator until all the DPBS evaporated. Pre-differentiated cells were dissociated and centrifuged as above, and pelleted cells were resuspended in Maturation Medium containing the following: 50% Neurobasal-A medium (GIBCO/Thermo Fisher Scientific; Cat. No. 10888022) and 50% DMEM/F-12 (GIBCO/Thermo Fisher Scientific; Cat. No. 11320033) as the base, 1X MEM Non-Essential Amino Acids, 0.5X GlutaMAX Supplement (GIBCO/Thermo Fisher Scientific; Cat. No. 35050061), 0.5X N-2 Supplement, 0.5X B-27 Supplement (GIBCO/Thermo Fisher Scientific; Cat. No. 17504044), 10 ng/mL NT-3, 10 ng/mL BDNF, 1 ug/mL Laminin mouse protein, and 2 ug/mL doxycycline hyclate. Pre-differentiated cells were subsequently counted and plated at 400,000-450,000 cells per well of a 12-well plate coated with Poly-L-Ornithine in 1 mL of Maturation medium with 20 uM trimethoprim (TMP) (Sigma-Aldrich, Cat No. 92131) to activate the CRISPRa machinery in these cells (TMP stabilizes the degron-tagged CRISPRa machinery). On day 7, half of the medium was removed and an equal volume of fresh Maturation medium without doxycycline was added. On day 14, half of the medium was removed and twice that volume of fresh medium without doxycycline was added. On day 19, neurons were harvested for sc-RNA-seq.

### Cell line generation and CRISPRa validation

#### K562 cells

K562 cells expressing dCas9-VP64 were generated in-house via lentiviral integration of a dCas9-VP64-blast construct^7^ (Addgene Plasmid #61422) into K562 cells. Cells were selected with 10 ug/mL blasticidin, and polyclonal cells were single-cell sorted into 96-well plates to grow up clonal cell lines expressing dCas9-VP64. Clonal cell lines were tested for CRISPRa activity by testing the ability of a CRISPRa gRNA to activate a minP-tdTomato construct^21^, and the highest tdTomato expressing cell line was used for experiments. K562 cells expressing dCas9-VPR were purchased from Horizon Discovery/Perkin Elmer (catalog ID: HD dCas9-VPR-005), and these cells were tested for CRISPRa activity using the same tdTomato expression assay described above.

#### iPSC-derived neurons

Human WTC11 iPSCs equipped with a doxycycline-inducible *NGN2* transgene expressed from the *AAVS1* safe-harbor locus as well as an ecDHFR-dCas9-VPH construct expressed from the CLYBL safe-harbor locus were a gift from the Kampmann lab^6^. These cells were tested for CRISPRa activity using the same tdTomato expression assay that was used to validate the K562 cell lines, which is described above.

### gRNA selection and design

A complete breakdown of gRNA library contents and overview of the gRNA design pipeline is illustrated in **Figure S1**. Briefly, enhancer-targeting gRNAs were selected from our CRISPRi library^2,46^. Specifically, 50 spacer sequences (2 per candidate enhancer) were randomly selected from the list of 664 significant “hit” enhancer-gene pairs in the at-scale library. Another 50 spacer sequences targeting an additional 25 candidate enhancers (again 2 per candidate enhancer) were randomly selected from candidate enhancer non-hits (*i.e.*, gRNAs from the at-scale library targeting candidate enhancer regions with strong biochemical marks predictive of regulatory activity that did not yield significant downregulation of any neighboring genes in our previous CRISPRi study). An additional 30 TSS-positive control gRNAs were randomly sampled from the top quartile of gRNAs recommended by Horlbeck *et al*. (hCRISPRa-v2 library)^16^. 50 NTC negative control spacer sequences were also selected from the hCRISPRa-v2 library^16^. The 313 candidate promoter targeting gRNAs were either selected from the Horlbeck *et al.* library^16^ or designed using FlashFry^46^ (**Figure S1**). Briefly, 50 candidate promoters of 9 NDD risk genes (*TCF4*, *FOXP1*, *SCN2A*, *CHD8*, *BCL11A*, *TBR1*, *SHANK3*, *SYNGAP1*, *ANK2*)^24,25^ were pulled from Basic GENCODE annotations^47^ and were filtered for “type” == “transcript” and “transcript_type” == “protein coding”. Separate bed files were generated for all promoter regions defined as the 500bp upstream of each protein coding transcript. Careful attention was paid to the strand orientation of each transcript when annotating promoter regions. Bed files were sorted and merged to combine multiple promoters with >1bp overlap into a single promoter annotation. Transcript bounds provided for each merged promoter begin +1bp from the end of the promoter and end at the position corresponding to the longest transcript mapping to that promoter. NGG-protospacer within these candidate promoters were identified using FlashFry and subsequently scored using default parameters (see FlashFry manuscript and user guide for a complete description of scoring metrics/algorithms)^46^. A TSS-distance metric was then calculated for each gRNA using human fetal brain 5’ Capped Analysis of Gene Expression (CAGE) data^48,49^ obtained from FANTOM (https://fantom.gsc.riken.jp/5/sstar/FF:10085-102B4; CTSS, hg38). First, the strongest FANTOM annotated TSS was identified within each +/-500 bp region up and downstream of each hg38 Gencode Basic protein coding transcript TSS. For regions with a tie between the highest scoring FANTOM TSSs, the TSS position closest to Gencode annotated TSS position was prioritized. Each candidate sgRNA from FlashFry was annotated with the distance to the nearest FANTOM TSS using the command “bedtools closest -a sgRNAs_with_fantom_tss -b strongest_fantom_tss_within_gencode_promoter -D b -t first.” For Gencode Basic protein coding transcripts without a human fetal brain FANTOM peak within 500 +/-bp, the distance of each sgRNA to the nearest Gencode TSS was reported instead. A distance of zero indicates that an sgRNA overlaps with the nearest annotated TSS. Multiple rounds of successively relaxing score and distance thresholds were then iterated until the top 4 gRNAs for each candidate promoter were selected (five selection rounds in total). Optimal TSS-distances were approximated using genome-wide CRISPRa design rules^36^. gRNAs flagged for potentially problematic polythymidine tracks or GC content were excluded. The gRNA selection criteria used in each round were as follows:

**Round 1: 1.** TSS Distance between −150 and −75 BP **2.** Doench2014OnTarget >= 0.2 **3.** Dangerous_in_genome <= 1 **4.** Hsu2013 > 80.
**Round 2: 1.** TSS Distance between −400 and −50 BP **2.** Doench2014OnTarget >= 0.2 **3.** Dangerous_in_genome <= 1 **4.** Hsu2013 > 80.
**Round 3: 1.** TSS Distance between −400 and −50 BP **2.** Doench2014OnTarget >= 0.2 **3.** Dangerous_in_genome <= 1 **4.** Hsu2013 > 50.
**Round 4: 1.** TSS Distance between −400 and −50 BP 2. Doench2014OnTarget >= 0.2 3. Dangerous_in_genome <= 2 4. Hsu2013 > 50.
**Round 5: 1.** Doench2014OnTarget >= 0.2 **2.** Dangerous_in_genome <= 2 **3.** Hsu2013 > 10 **4.** DoenchCFD_maxOT < 0.95

Complete oligo sequences with gRNA spacers and additional sequences for cloning into piggyFlex are listed in **Table S1**. Note all gRNAs in our library are designed/modified to start with a G followed by the 19 base pair spacer to facilitate Pol III transcription.

### gRNA library cloning into piggyFlex vector

The 493 gRNAs with associated 10N random barcodes were ordered as an IDT oPool and PCR amplified with Q5 High-Fidelity polymerase (NEB, Cat. No. M0491S) to make double stranded DNA. The piggyFlex backbone vector was digested with SalI (NEB, Cat. No. R3138S) and BbsI (NEB, Cat. No. R0539S) in 10X NEBuffer r2 at 37 degrees Celsius overnight to ensure complete digestion of the backbone. This digestion cuts out the EF1a-puro-GFP cassette of the vector which is then added back in a later cloning step. The digestion product was run on a 1% agarose gel in TAE buffer, and the linear backbone vector (5098 base pairs in size) was gel extracted using a gel extraction kit (NEB, Cat. No. T1020S). The second product from the digestion (2878 base pairs) which contains the EF1a-puro-GFP cassette was saved for a later assembly reaction in the final cloning step (described below). The PCR amplified IDT oPool gRNAs with associated 10N random barcodes were cloned into the linear backbone using NEBuilder HiFi DNA Assembly (NEB, Cat. No. E2621S) using 0.15 pmol of the insert (gRNA library) and 0.02 pmol of the linear backbone. Assembled product was transformed into electrocompetent cells (NEB, Cat. No. C3020K) and plasmid DNA was extracted with a midiprep kit (Zymo Research, Cat. No. D4200). The resulting vector was then digested with SapI (NEB, Cat. No. R0569S), for one hour at 37 degrees Celsius. Digested product was cleaned with 0.5X AMPure beads (Beckman Coulter, Cat. No. A63880) and cleaned digested linear backbone was used for a subsequent assembly reaction to add the EF1a-puro-GFP cassette back into the final piggyFlex vector between the gRNA sequence and the 10N random barcode sequences. 0.014 pmol of the linear backbone was assembled with 0.056 pmol of the insert sequence and the assembly reaction was cleaned with a 0.5X AMPure step. The assembled product was transformed into electrocompetent cells and plasmid DNA was extracted with a midiprep kit. The final plasmid library was subsequently PCR amplified and sequenced to ensure that all 493 gRNAs were successfully cloned into the piggyFlex vector. Note: The 10N barcode is an additional gRNA identification strategy that can be used to assign gRNAs to cells, however, we used directly sequenced gRNAs (from the 10x Genomics capture sequence) to identify gRNAs in this work as this more accurately assigns gRNA transcripts to cells^40^.

### Transfection of the gRNA library, selection, and cell culture

#### K562 cells

16 million K562 cells (8 million K562-VP64 cells and 8 million K562-VPR cells) were transfected with the gRNA library and the piggyBac transposase (System Biosciences, Cat. No. PB210PA-1) at a 20:1 molar ratio of library:transposase using a Lonza 4D nucleofector and the Lonza nucleofector protocol for K562 cells. The 16 million cells were split across 8 100 uL nucleofection cartridges, with each individual nucleofection cartridge receiving 2 million cells and 2 ug of total DNA. After nucleofection, cells were transferred to pre-warmed RPMI media in a cell culture flask and incubated at 37 degrees Celsius. One day after transfection, cells were selected with 2 ug/mL puromycin (GIBCO/Thermo Fisher Scientific; Cat. No. A1113803). After 9 days, cells were harvested for single-cell transcriptome profiling.

#### Induced pluripotent stem cells

6 million dCas9-VPH iPSCs (same cells as described above) were transfected with the gRNA library and the piggyBac transposase at a 5:1 molar ratio of library:transposase using the Lonza nucleofector and the Lonza nucleofector CB-150 program. The 6 million cells were split across 6 100 uL nucleofection cartridges, with each individual nucleofection cartridge receiving 1 million cells and 17.5 ug of total DNA. After nucleofection, cells were transferred to pre-warmed mTeSr Plus basaI medium with ROCK inhibitor in a cell culture flask and incubated at 37 degrees Celsius. One day after transfection, cells were selected with 20 ug/mL puromycin (note: the AAVS1-NGN2 construct has a puromycin resistance cassette on it, so a higher dose of puromycin was used to successfully select for cells that received a gRNA in the presence of an existing puromycin resistance cassette). Media changes were performed daily (mTeSr Plus basaI medium with ROCK inhibitor and 10 ug/mL puromycin) for seven days prior to initiating neuron differentiation (described in “Human iPSC-derived neuronal cell culture, differentiation, and CRISPRa induction” methods section).

### 10x Genomics sc-RNA-seq with associated gRNA transcript capture

#### K562 screen

Cells were harvested and prepared into single-cell suspensions following the 10x Genomics Single Cell Protocols Cell Preparation Guide (Manual part number CG00053, Rev C). Four lanes were used for the single-cell transcriptome profiling, with two lanes containing cells from the K562-VP64 cell line, and two lanes containing cells from the K562-VPR cell line. Roughly 10,000 cells were captured per lane of a 10x Chromium chip (Next GEM Chip G) using Chromium Next GEM Single Cell 3ʹ Reagent Kits v3.1 with Feature Barcoding technology for CRISPR Screening (10x Genomics, Inc, Document number CG000205, Rev D).

#### iPSC-derived neuron screen

iPSC-derived neurons were harvested and prepared into single-cell suspensions following a published protocol^50^. Cells were split into two batches, with one batch going through a fluorescence-activated cell sorting (FACS) step to sort on the top 40% of green fluorescent protein (GFP) expression to enrich for neurons with greater numbers of gRNAs integrated, and the second batch going directly into the 10x Genomics single-cell library preparation protocol. Sorting on the top 40% of GFP expression resulted in a two-fold increase in the mean number of gRNAs integrated in those cells as compared to unsorted cells. Four lanes were used for the single-cell transcriptome profiling, with two lanes containing GFP-positive sorted cells, and two lanes containing unsorted cells. Roughly 13,000 cells were captured per lane of a 10x Chromium high-throughput chip (Next GEM Chip M) using Chromium Next GEM Single Cell 3’ HT Reagent Kits v3.1 (Dual Index) with Feature Barcode technology for CRISPR Screening (10x Genomics, Inc, Document number CG000418, Rev C).

#### Sequencing of scRNA-seq libraries

Final libraries were sequenced on an Illumina NextSeq 2000 P3 100 cycle kit (R1:28 I1:10, I2:10, R2:90) for each screen (K562 and iPSC-derived neuron screens). Gene expression and gRNA transcript libraries were pooled at a 4:1 ratio for sequencing.

#### Transcriptome data processing and quality control filtering for K562 and iPSC-derived neuron screens

CellRanger version 6.0.1 was used to perform bcl2fastq and count matrix generation. CellRanger mkfastq was run using default parameters, and CellRanger count was run using the GRCh38-3.0.0 reference transcriptome from 10x Genomics and default parameters. For the K562 screen, cells with greater than 10% mitochondrial reads and less than 4000 UMIs were filtered out. For the iPSC-derived neuron screen, cells with greater than 17% mitochondrial reads and less than 1500 unique molecular identifiers (UMIs) were filtered out. After quality control filtering, 33,944 cells were retained in the K562 screen, and 51,183 cells were retained in the iPSC-derived neuron screen. The resulting count matrix output after this filtering was used for all downstream analyses.

#### Neuron differentiation transcriptome projection

Single-cell transcriptome data from a time course study of iPSC-derived neurons^27^ was downloaded from https://www.ebi.ac.uk/biostudies/arrayexpress/studies/E-MTAB-10632 (Accession No. E-MTAB-10632, matrices_timecourse.tar.gz), and integrated with the neuron CRISPRa screening dataset described here. Seurat v4 was used for all data analyses^51^. The CRISPRa dataset was randomly downsampled to 5,000 cells for this analysis. Count matrices from both matrices were filtered to include only shared genes from the two datasets (n=14,777 genes). SelectIntegrationFeatures() and FindIntegrationAnchors() were run using default parameters to identify anchors for integration. 20,606 anchors were found and 2,953 anchors were retained for data integration. IntegrateData() was run using the retained 2,953 anchors to integrate the two datasets. After integration, standard Seurat single-cell analysis was performed to scale the data, and run the PCA and UMAP algorithms.

### gRNA assignment and differential gene expression testing

Genomic coordinates (hg38) for final gRNA spacers were isolated using a loop built around the matchPattern() function from the BSgenome package^52^. A 2Mb window (1Mb upstream and downstream) around each gRNA was then calculated and all genes within the 2Mb window were isolated using a loop built around ENSEMBL biomaRt getBM() function^53,54^. These 1Mb neighboring gene sets were then filtered to unique entries (unique HGNC symbols) for compatibility with the Seurat FindMarkers() function used in DE testing.

A global UMI filter of 5 gRNA UMIs/cell was used to assign gRNAs to single cell transcriptomes for both K562 and iPSC-derived neuron datasets (note this heuristic threshold was chosen based on manual inspection of the UMI count distributions for each gRNA and prior work)^2^. gRNA UMI counts for each cell were derived from the count matrix of passing cells output by CellRanger (which applies an automatic total UMI threshold to cells) and which also passed QC.

In our original approach, expression of a given gene within 1Mb of the gRNA of interest was compared between all cells with a given gRNA and all other cells as control. log2() fold changes in expression for a given gene were calculated using the Seurat FindMarkers() function with the following arguments: ident.1 = gRNA_Cells, ident.2 = Control_Cells, min.pct = 0, min.cells.feature = 0, min.cells.group = 0, features = target_gene, logfc.threshold = 0. A Wilcoxon rank-sum test was used to generate raw differential expression *P*-values. This process was then iterated for all genes within 1Mb of all gRNAs. NTCs were tested against all genes within 1Mb of any targeting gRNA. Only tests involving genes detected in >0.2% of test gRNA and control cells were carried forward.

These raw differential expression *P*-values were then used to calculate empirical *P*-values to call EFDR < 0.1 sets^2^. Specifically, an empirical *P*-value was calculated for each gRNA-gene test as:

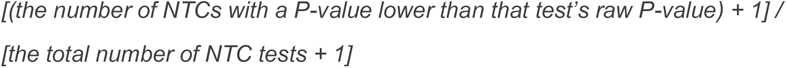

Empirical *P*-values were then Benjamini-Hochberg corrected, and those < 0.1 were kept for 10% EFDR sets.

With SCEPTRE, count matrices and associated metadata were used to construct a single cell covariate matrix with gRNA and gene expression library size (total UMIs), unique genes (non-zero expression in gRNA and gene expression libraries), cell line (in the case of K562 cells with differing CRISPRa effectors), whether cells were GFP sorted prior to profiling, 10X lane, and percent mitochondrial reads input as covariates during model fitting. A global UMI filter of 5 gRNA UMIs per cell was used to assign gRNAs to single cell transcriptomes. The run_sceptre_highmoi_experimental() function was used to run calibration (NTC) and discovery (tests) with default parameters (e.g. two-tailed, >7 targeting and control cells with non-zero expression). Each individual gRNA was tested against all neighboring genes within 1Mb. NTCs were tested against all genes within 1Mb of a targeting gRNA. NTCs were randomly downsampled to match the number of targeting cis tests for visualization on QQ plots. Resulting SCEPTRE *P*-values were then Benjamini-Hochberg corrected and those < 0.1 were kept for two-sided discovery sets.

Log2 fold changes between gRNA and control cells were visualized using the gviz package^55^ along with tracks for RefSeq transcripts (ENSEMBL biomaRt), H3K27ac, and ATAC seq peaks. The K562 ATAC and H3K27ac data were downloaded from ENCODE ^56^. ATAC-seq and H3K27ac CUT&RUN data from 7-8 week old NGN2-iPSC inducible excitatory neurons was obtained from Song et al. 2019^45^. As previously described, ATAC-seq and CUT&RUN reads were trimmed to 50bp using TrimGalore with the command --hardtrim5 50 before alignment (https://github.com/FelixKrueger/TrimGalore). ATAC-seq reads were realigned to hg38 using the standard Encode Consortium ATAC-seq and ChIP-seq pipelines respectively with default settings and pseudo replicate generation turned off. Trimmed, sorted, duplicate and chrM removed ATAC-seq bam files from multiple biological replicates were combined into a single bam file using samtools merge v1.10^57^. Trimmed CUT&RUN reads were realigned to hg38 using Bowtie2 v2.3.5.1 with the following settings --local --very-sensitive-local --no-mixed --no-discordant -I 10 -X 700 and output sam files were convert to bam format using samtools view^57,58^. Duplicated reads were removed from the CUT&RUN bam file using Picard MarkDuplicates v2.26.0 with the -- REMOVE_DUPLICATES =true and --ASSUME_SORTED=true options (http://broadinstitute.github.io/picard/). Finally, bam files were converted using the bedtools genomecov followed by the UCSC bedGraphToBigWig utility.

For correlations of epigenetic features with CRISPRa hits, epigenetic feature datasets were downloaded from ENCODE or corresponding primary publications (datasets listed in **Table S8**). Whether a gRNA was a SCEPTRE hit (Benjamini-Hochberg–adjusted SCEPTRE *P-*value < 0.1) was treated as a categorical input variable for Spearman correlations with epigenetic features quantified in 100bp windows at the gRNA target sites. Similarly, various gRNA metrics (e.g. GC content) or gene level metrics (e.g. baseline expression) were previously quantified, either as part of the gRNA design process or gRNA assignment and differential expression testing frameworks (described above) and tested for association with hit gRNAs.

### Singleton replication and validations with bulk RNA-sequencing

To replicate and validate a gRNA’s ability to upregulate its target gene outside of the pooled screening format, we generated individual, polyclonal cell lines that each expressed a single gRNA from a chosen representative set of gRNAs (**Table S15**). This representative gRNA set was chosen to validate gRNAs that upregulate their targets in a cell-type specific manner (TBR1 is neuron-specific; ASIC1 and ANXA1 are enhancer targeting gRNAs that are K562-specific) and to validate promoter targeting gRNAs that are able to upregulate their targets in both cell types (e.g. BCL11A, TCF4, and FOXP1). gRNAs were ordered in two Oligo Pools (Integrated DNA Technologies), with each gRNA bearing a unique handle sequence that was used to uniquely PCR amplify out individual gRNA oligos (**Table S15**). gRNAs were individually cloned into the piggyFlex vector exactly as described in the “gRNA library cloning into piggyFlex vector” methods section above, with the only difference being that the 10N barcode was not included in the oligo design for these validation oligos. After cloning, the gRNA sequences in the piggyFlex vectors were sequence verified to ensure correct insertion of the gRNA sequences.

PiggyFlex constructs containing gRNA sequences were transfected into K562 cells and iPSCs with the Lonza 4D nucleofector (using the K562 protocol for K562 cells, and the CB-150 program for the iPSCs). 525 ng of total DNA (using a 5:1 molar ratio of piggyFlex vector and transposase vector) was transfected into three replicates of 30,000 cells each. After transfection, cells were seeded into 48-well plates for culture. One day later, 2 ug/mL (for K562 cells) and 20 ug/mL (for iPSCs) puromycin (GIBCO/Thermo Fisher Scientific; Cat. No. A1113803) was added to the cultures to select for successfully transfected cells. Daily media changes were performed (with puromycin) to continually select cells for seven days. After seven days of selection, K562 cells were lysed with TRIzol (Invitrogen, Cat. No. 15596026). Samples were frozen in TRIzol reagent at −20C, and RNA was extracted using the Direct-zol RNA miniprep kit (Zymo Research, Cat. No. R2050). After puromycin selection, iPSCs were differentiated into neurons exactly as described in the “Human iPSC-derived neuronal cell culture, differentiation, and CRISPRa induction” section. iPSC-derived neurons were differentiated for eight days prior to TRIzol lysis and RNA extraction as described above for K562 cells.

RNA-sequencing libraries were prepared using an in-house library preparation method. 30-50 ng of RNA was used as input for each library. RNA was reverse transcribed using indexed oligo dT primers (ACGACGCTCTTCCGATCTNNNNNNNNAGAGAACTTGTTTTTTTTTTTTTTTTTTTTTTTTTTTTTTVN), template switching oligo (AAGCAGTGGTATCAACGCAGAGTGAATGGG), and Template Switching RT Enzyme Mix protocol from NEB (New Engand Biolabs, Cat. No. #M0466L). cDNA was PCR amplified using the NEBNext High-Fidelity 2X PCR Master Mix (New England Biolabs, Cat. No. M0541) with the following forward and reverse primers: ACGACGCTCTTCCGATC (forward), AAGCAGTGGTATCAACGCA (reverse). SYBR Green (Thermo Fisher Scientific, Cat. No. S7567) was added to track the amplification curve and proper PCR cycle number. PCR amplified cDNA was purified with 1X AMPure beads (Beckman Coulter, Cat. No. A63882), washed twice with 70% ethanol, eluted in 20 uL of nuclease-free water and loaded on an Agilent Tapestation using D5000 reagents (Agilent Technologies, Cat. No. 5067-5589) and the corresponding screentape (Agilent Technologies, Cat. No. 5067-5588). 30-50 ng of purified PCR amplified cDNA was fragmented using an i7 loaded Tn5 transposase similar to the process described by Cao *et al*.^43^. To load the transposase, we followed the manufacturer’s protocol to load unloaded transposase (Diagenode, Cat. No. C01070010-20). Tagmented DNA was PCR amplified using NEBNext High-Fidelity 2X PCR Master Mix (New England Biolabs, Cat. No. M0541) with indexed primers (CAAGCAGAAGACGGCATACGAGATNNNNNNNNNNGTCTCGTGGGCTCGG and AATGATACGGCGACCACCGAGATCTACACNNNNNNNNNNACACTCTTTCCCTACACGACGCTCTTCCGATCT) following previously described PCR conditions by Cao *et al.*^43^. This PCR adds sample-specific indices and P5 and P7 Illumina sequencing adapters. Libraries were cleaned using a 1X AMPure bead cleanup (Beckman Coulter, Cat. No. A63882), and quantified on an Agilent TapeStation using D1000 reagents (Agilent, Cat. No. 5067-5583) and the corresponding screentape (Agilent, Cat. No. 5067-5582). For sequencing, libraries were diluted to 2nM and pooled equimolarly, and then diluted to a loading concentration of 650 pM. Libraries were sequenced on an Illumina NextSeq 2000 using P1 100 cycle kits (R1: 18, I1:10, I2: 10 (optional), and R2:110). Transcript and gene-level quantifications were performed using kallisto^59^. Prior to transcript quantification, reads were downsampled to the minimum number of reads observed for a sample to control for differences in sequencing depth. DESeq2^60^ was used to quantify expression changes for all genes within 1Mb of the gRNA target site and genome wide. Genome-wide differential analyses were conducted using default parameters. Target gene expression was compared between gRNA cells and control cells using a two-tailed Wilcoxon rank-sum test with a significance threshold of 0.1 (mirroring single-cell analysis framework described above). Gviz was used to visualize differential expression testing results alongside tracks for neighboring genes (ENSEMBL biomaRt) as described above.

## References

1. Gasperini, M., Tome, J. M. & Shendure, J. Towards a comprehensive catalogue of validated and target-linked human enhancers. Nat. Rev. Genet. 21, 292–310 (2020).

2. Gasperini, M. et al. A Genome-wide Framework for Mapping Gene Regulation via Cellular Genetic Screens. Cell 176, 377–390.e19 (2019).

3. Fulco, C. P. et al. Systematic mapping of functional enhancer–promoter connections with CRISPR interference. Science 354, 769–773 (2016).

4. Xie, S., Duan, J., Li, B., Zhou, P. & Hon, G. C. Multiplexed Engineering and Analysis of Combinatorial Enhancer Activity in Single Cells. Mol. Cell 66, 285–299.e5 (2017).

5. Gilbert, L. A. et al. Genome-Scale CRISPR-Mediated Control of Gene Repression and Activation. Cell 159, 647– 661 (2014).

6. Tian, R. et al. Genome-wide CRISPRi/a screens in human neurons link lysosomal failure to ferroptosis. Nat. Neurosci. 24, 1020–1034 (2021).

7. Konermann, S. et al. Genome-scale transcriptional activation by an engineered CRISPR-Cas9 complex. Nature 517, 583–588 (2014).

8. Schmidt, R. et al. CRISPR activation and interference screens decode stimulation responses in primary human T cells. Science 375, eabj4008 (2022).

9. Simeonov, D. R. et al. Discovery of stimulation-responsive immune enhancers with CRISPR activation. Nature 549, 111–115 (2017).

10. Matharu, N. et al. CRISPR-mediated activation of a promoter or enhancer rescues obesity caused by haploinsufficiency. Science 363, (2019).

11. Tamura, S. et al. CRISPR activation rescues abnormalities in SCN2A haploinsufficiency-associated autism spectrum disorder. bioRxiv 2022.03.30.486483 (2022) doi:10.1101/2022.03.30.486483.

12. Dai, Z. et al. Inducible CRISPRa screen identifies putative enhancers. J. Genet. Genomics 48, 917–927 (2021).

13. Tak, Y. E. et al. Augmenting and directing long-range CRISPR-mediated activation in human cells. Nat. Methods 18, 1075–1081 (2021).

14. Joung, J. et al. Genome-scale activation screen identifies a lncRNA locus regulating a gene neighbourhood. Nature 548, 343–346 (2017).

15. Zhou, B. et al. Comprehensive, integrated, and phased whole-genome analysis of the primary ENCODE cell line K562. Genome Res. 29, 472–484 (2019).

16. Horlbeck, M. A. et al. Compact and highly active next-generation libraries for CRISPR-mediated gene repression and activation. Elife 5, (2016).

17. Matharu, N. & Ahituv, N. Modulating gene regulation to treat genetic disorders. Nat. Rev. Drug Discov. 19, 757–775 (2020).

18 Lalanne, J.-B. et al. Multiplex profiling of developmental enhancers with quantitative, single-cell expression reporters. bioRxiv 2022.12.10.519236 (2022) doi:10.1101/2022.12.10.519236.

19. Hill, A. J. et al. On the design of CRISPR-based single-cell molecular screens. Nat. Methods 15, 271–274 (2018).

20. Maeder, M. L. et al. CRISPR RNA–guided activation of endogenous human genes. Nat. Methods 10, 977–979 (2013).

21. Chavez, A. et al. Highly efficient Cas9-mediated transcriptional programming. Nat. Methods 12, 326–328 (2015).

22. Esvelt, K. M. et al. Orthogonal Cas9 proteins for RNA-guided gene regulation and editing. Nat. Methods 10, 1116– 1121 (2013).

23. Dixon, J. R. et al. Topological domains in mammalian genomes identified by analysis of chromatin interactions. Nature vol. 485 376–380 Preprint at 10.1038/nature11082 (2012).

24. Satterstrom, F. K. et al. Large-Scale Exome Sequencing Study Implicates Both Developmental and Functional Changes in the Neurobiology of Autism. Cell 180, 568–584.e23 (2020).

25. Fu, J. M. et al. Rare coding variation provides insight into the genetic architecture and phenotypic context of autism. Nat. Genet. 54, 1320–1331 (2022).

26. Zhang, Y. et al. Rapid single-step induction of functional neurons from human pluripotent stem cells. Neuron 78, 785–798 (2013).

27. Wang, C. et al. Scalable Production of iPSC-Derived Human Neurons to Identify Tau-Lowering Compounds by High-Content Screening. Stem Cell Reports 9, 1221–1233 (2017).

28. Lin, H.-C. et al. NGN2 induces diverse neuron types from human pluripotency. Stem Cell Reports 16, 2118–2127 (2021).

29. Barry, T., Wang, X., Morris, J. A., Roeder, K. & Katsevich, E. SCEPTRE improves calibration and sensitivity in single-cell CRISPR screen analysis. Genome Biol. 22, 344 (2021).

30. Morris, J. A. et al. Discovery of target genes and pathways at GWAS loci by pooled single-cell CRISPR screens. Science 380, eadh7699 (2023).

31. Barry, T., Mason, K., Roeder, K. & Katsevich, E. Robust differential expression testing for single-cell CRISPR screens at low multiplicity of infection. bioRxiv 2023.05.15.540875 (2023) doi:10.1101/2023.05.15.540875.

32. Barry, T., Roeder, K. & Katsevich, E. Exponential family measurement error models for single-cell CRISPR screens. arXiv [stat.ME] (2022).

33. Tamura, S. et al. CRISPR activation rescues abnormalities in SCN2A haploinsufficiency-associated autism spectrum disorder. bioRxiv 2022.03.30.486483 (2022) doi:10.1101/2022.03.30.486483.

34. Matharu, N. & Ahituv, N. Modulating gene regulation to treat genetic disorders. Nat. Rev. Drug Discov. 19, 757–775 (2020).

35. Matharu, N. et al. CRISPR-mediated activation of a promoter or enhancer rescues obesity caused by haploinsufficiency. Science 363, (2019).

36. Sanson, K. R. et al. Optimized libraries for CRISPR-Cas9 genetic screens with multiple modalities. Nat. Commun. 9, 5416 (2018).

37. Yao, D. et al. Multi-center integrated analysis of non-coding CRISPR screens. bioRxiv 2022.12.21.520137 (2022) doi:10.1101/2022.12.21.520137.

38. Tycko, J. et al. Development of compact transcriptional effectors using high-throughput measurements in diverse contexts. bioRxiv 2023.05.12.540558 (2023) doi:10.1101/2023.05.12.540558.

39. Murphy, A. E., Fancy, N. N. & Skene, N. G. Avoiding false discoveries: Revisiting an Alzheimer’s disease snRNA-Seq dataset. bioRxiv 2023.04.01.535040 (2023) doi:10.1101/2023.04.01.535040.

40. Replogle, J. M. et al. Combinatorial single-cell CRISPR screens by direct guide RNA capture and targeted sequencing. Nat. Biotechnol. 38, 954–961 (2020).

41. Marshall, J. L. et al. HyPR-seq: Single-cell quantification of chosen RNAs via hybridization and sequencing of DNA probes. Proc. Natl. Acad. Sci. U. S. A. 117, 33404–33413 (2020).

42. Schraivogel, D. et al. Targeted Perturb-seq enables genome-scale genetic screens in single cells. Nat. Methods 17, 629–635 (2020).

43. Cao, J. et al. The single-cell transcriptional landscape of mammalian organogenesis. Nature 566, 496–502 (2019).

44. Xu, Z., Sziraki, A., Lee, J., Zhou, W. & Cao, J. PerturbSci-Kinetics: Dissecting key regulators of transcriptome kinetics through scalable single-cell RNA profiling of pooled CRISPR screens. bioRxiv 2023.01.29.526143 (2023) doi:10.1101/2023.01.29.526143.

45. Song, M. et al. Mapping cis-regulatory chromatin contacts in neural cells links neuropsychiatric disorder risk variants to target genes. Nat. Genet. 51, 1252–1262 (2019).

46. McKenna, A. & Shendure, J. FlashFry: a fast and flexible tool for large-scale CRISPR target design. BMC Biol. 16, 74 (2018).

47. Frankish, A. et al. GENCODE reference annotation for the human and mouse genomes. Nucleic Acids Res. 47, D766–D773 (2019).

48. Lizio, M. et al. Update of the FANTOM web resource: expansion to provide additional transcriptome atlases. Nucleic Acids Res. 47, D752–D758 (2019).

49. FANTOM Consortium and the RIKEN PMI and CLST (DGT), et al. A promoter-level mammalian expression atlas. Nature 507, 462–470 (2014).

50. Jerber, J., Haldane, J., Steer, J., Pearce, D. & Patel, M. Dissociation of neuronal culture to single cells for scRNA-seq (10x Genomics) v1. Preprint at 10.17504/protocols.io.bh32j8qe.

51. Hao, Y. et al. Integrated analysis of multimodal single-cell data. Cell 184, 3573–3587.e29 (2021).

52. Pagès, H. BSgenome: Software infrastructure for efficient representation of full genomes and their SNPs. R package version.

53. Durinck, S., Spellman, P. T., Birney, E. & Huber, W. Mapping identifiers for the integration of genomic datasets with the R/Bioconductor package biomaRt. Nat. Protoc. 4, 1184–1191 (2009).

54. Durinck, S. et al. BioMart and Bioconductor: a powerful link between biological databases and microarray data analysis. Bioinformatics 21, 3439–3440 (2005).

55. Hahne, F. & Ivanek, R. Visualizing Genomic Data Using Gviz and Bioconductor. Methods in Molecular Biology 335– 351 Preprint at 10.1007/978-1-4939-3578-9_16 (2016).

56. Rosenbloom, K. R. et al. ENCODE data in the UCSC Genome Browser: year 5 update. Nucleic Acids Res. 41, D56–63 (2013).

57. Li, H. et al. The Sequence Alignment/Map format and SAMtools. Bioinformatics 25, 2078–2079 (2009).

58. Langmead, B. & Salzberg, S. L. Fast gapped-read alignment with Bowtie 2. Nat. Methods 9, 357–359 (2012).

59. Bray, N. L., Pimentel, H., Melsted, P. & Pachter, L. Erratum: Near-optimal probabilistic RNA-seq quantification. Nat. Biotechnol. 34, 888 (2016).

60. Love, M. I., Huber, W. & Anders, S. Moderated estimation of fold change and dispersion for RNA-seq data with DESeq2. Genome Biol. 15, 550 (2014).

